# *Aspergillus fumigatus* can display persistence to the fungicidal drug voriconazole

**DOI:** 10.1101/2022.05.16.491816

**Authors:** Jennifer Scott, Clara Valero, Álvaro Mato-López, Ian J. Donaldson, Alejandra Roldán, Harry Chown, Norman Van Rhijn, Rebeca Lobo-Vega, Sara Gago, Takanori Furukawa, Alma Morogovsky, Ronen Ben Ami, Paul Bowyer, Nir Osherov, Thierry Fontaine, Gustavo H. Goldman, Emilia Mellado, Michael Bromley, Jorge Amich

**Author notes:** These authors contributed equally (Jennifer Scott, Clara Valero). They are ordered alphabetically.

## Abstract

*Aspergillus fumigatus* is a filamentous fungus that can infect the lungs of patients with immunosuppression and/or underlying lung-diseases. The mortality associated with chronic and invasive aspergillosis infections remain very high, despite availability of antifungal treatments. In the last decade, there has been a worrisome emergence and spread of resistance to the first line antifungals, the azoles. The mortality caused by resistant isolates is even higher, and patient management is complicated as the therapeutic options are reduced. Nevertheless, treatment failure is also common in patients infected with azole-susceptible isolates, which can be due to several non-mutually exclusive reasons, such as poor drug absorption. In addition, the phenomena of tolerance or persistence, where susceptible pathogens can survive the action of an antimicrobial for extended periods, have been associated with treatment failure in bacterial infections, and their occurrence in fungal infections already proposed. Here, we demonstrate that some isolates of *A. fumigatus* display persistence to voriconazole. A sub-population of the persister isolates can survive for extended periods and even grow at slow rates in the presence of supra-MIC (Minimum Inhibitory Concentration) of voriconazole and seemingly other azoles. Persistence cannot be eradicated with adjuvant drugs or antifungal combinations and seems to reduce the efficacy of treatment for certain individuals in a *Galleria mellonella* model of infection. Furthermore, persistence implies a distinct transcriptional profile, demonstrating that it is an active response. We propose that azole persistence might be a relevant and underestimated factor that could influence the outcome of infection in human aspergillosis.

**IMPORTANCE:** The phenomena of antibacterial tolerance and persistence, where pathogenic microbes can survive for extended periods in the presence of cidal drug concentrations, have received significant attention in the last decade. Several mechanisms of action have been elucidated, and their relevance for treatment failure in bacterial infections demonstrated. In contrast, our knowledge about antifungal tolerance and, particularly, persistence are still very scarce. In this study, we have characterised the response of the prominent fungal pathogen *Aspergillus fumigatus* to the first line therapy antifungal voriconazole. We comprehensively show that some isolates display persistence to this fungicidal antifungal and identify various potential mechanisms of action. In addition, using an alternative model of infection, we provide initial evidence to suggest that persistence may cause treatment failure in some individuals. Therefore, we propose that azole persistence is an important factor to consider and further investigate in *A. fumigatus*.

## INTRODUCTION

*Aspergillus fumigatus* is the most prominent fungal pathogen of the human lung, being the major causative agent of a range of diseases collectively termed aspergillosis [1]. The incidence of fatal aspergillosis infections is on the rise due to the increase in the at-risk population [2, 3], including severe COVID-19 patients [4–6]. The mortality associated with *A. fumigatus* infections remains unacceptably high: 38% 5-year mortality rates for chronic pulmonary aspergillosis [7] and ranging from ∼35% in early diagnosed and treated patients to nearly 100% if diagnosis is missed or delayed in invasive aspergillosis [8]. These number represent the highest mortality of all invasive fungal diseases [9]. This situation has become even more worrisome due to the emergence of clinical resistance to existing antifungals, which poses a serious threat to human health [10]. Azoles are currently the only FDA-approved class of mould-active agents that can be administrated orally and intravenously, and accordingly they are used for both the evidence-based treatment and prevention of aspergillosis diseases [11]. However, in the last decade, the number of clinical *A. fumigatus* isolates that are resistant to triazole drugs has increased worldwide, causing serious problems for clinical management, as the therapeutic options are reduced and the mortality rates caused by resistant isolates are higher [7, 12, 13]. Furthermore, both in clinical disease and in experimental animal models, it has been observed that treatment failure is common even if the infecting fungal isolate is susceptible to the azole used for treatment [14, 15]. There are several possible explanations for treatment failure in these cases, including antifungal tolerance or persistence [16, 17].

In the last decade there has been a revolution in our understanding of how pathogenic microorganisms withstand the challenge of antimicrobial drugs. Classically, pathogens were described to be either susceptible or resistant to a certain drug. Antimicrobial resistance is due to a genetic feature in the microbe (intrinsic or acquired *via* mutations) that enables it to grow normally at high concentrations of the agent, above the Minimum Inhibitory Concentration (MIC) defined for the antimicrobial based on the clinical breakpoint. However, in recent years it has become obvious that microbes can withstand the action of drugs by at least three other mechanisms: tolerance, persistence, or heteroresistance. These three phenomena share the feature that they are not triggered by mutations in the microbe’s DNA sequence, yet differ in other details. Drug tolerance has been defined as the ability of all cells of a genetically isogenic strain to survive, and even grow at slow rates, for extended periods in the presence of drug concentrations that are greater than the minimal inhibitory concentration (MIC). In persistence, only a small fraction of the isogenic population (usually <1% of the cells) survive, and grow at slow rates, for an extended period of time in the presence of supra-MIC concentrations of the drug [18, 19]. These two phenomena do not imply an increase in the MIC. Finally, heteroresistance is the transient capacity of a microbial subpopulation to increase the MIC in the presence of the drug, due to adaptation, epigenetic modifications or reversible aneuploidy. All those phenomena have been implicated in treatment failure and relapse in bacterial infections [18, 20]. These definitions are summarised in Table 1.

**Table 1.** Current and proposed definitions for the phenomena of resistance, heteroresistance, tolerance and persistence in pathogenic fungi (see discussion).

| Phenomenon | Previous (bacteria) Definition | Adjustment for Fungi |
| --- | --- | --- |
| **Resistance** | Stable capacity of an isolate to grow normally in the presence of high concentrations of the drug, above the minimum inhibitory concentration (MIC) defined for the antimicrobial based on the clinical breakpoint. This capacity is based on specific genetic sequence(s) (intrinsic or acquired) in the genome. | The same definition for fungi |
| **Heteroresistance** | Transient capacity of a subpopulation of cells to increase the MIC in the presence of the drug, without a modification in the genome sequence. | The same definition for fungi |
| **Tolerance** | Ability of <u>all cells</u> of a susceptible, genetically isogenic strain to survive, and even grow at slow rates, for extended periods in the presence of drug concentrations that are greater than the MIC. | In the presence of <u>fungistatic drugs</u>, capacity of all or high proportion of the cells of a susceptible isolate to grow in the presence of drug concentrations that are greater than the MIC. |
| **Persistence** | Ability of a <u>small subpopulation of cells</u> of a susceptible, genetically isogenic strain to survive, and even grow at slow rates, for extended periods in the presence of drug concentrations that are greater than the MIC. | In the presence of <u>fungicidal drugs</u>, capacity of a small fraction of cells of a susceptible isolate to survive, and even grow at slow rates, for extended periods in the presence of drug concentrations that are greater than the MIC. |

In fungi, tolerance has been often described as “trailing growth” and its relevance for infection has been mostly disregarded. However, a recent landmark study in *Candida albicans* has characterised tolerance as a distinct, strain-specific feature and provided evidence for its relevance in persistent candidemia [17, 21]. In *Aspergillu*s, trailing growth is known to be prominent in the presence of caspofungin [22, 23] apparently due to single strain heterogeneity [24]. Moreover, at higher concentrations of this drug some isolates can resume normal growth, an effect known as “Eagle” or paradoxical effect (see review [25] for more information), which we have recently demonstrated as a tolerance phenotype [26]. The possible relevance of the paradoxical effect in clinical practice is still under debate, but there is evidence suggesting that it might be important [27], and concern has been raised by some clinicians [28]. Remarkably, it seems that the phenomenon of tolerance in fungi is seen with static drugs, e.g. azoles for *C. albicans* and echinocandins for *A. fumigatus*. However, the possibility that *A. fumigatus* can display tolerance or persistence to azole antifungals had not been previously investigated. Interestingly, in contrast to *C. albicans* and *Cryptococcus neoformans*, azoles have been shown to have fungicidal activity against *A. fumigatus* [29–31], which implies that the underlying mechanisms are likely different. Whilst these phenomena have so far only been investigated in bacteria or yeasts, the approaches to detect and investigate azole tolerance or persistence need to be tailored in filamentous fungi, like *A. fumigatus*, as these organisms form multicellular hyphae. Using various complementary approaches, here we show that small subpopulations of certain *A. fumigatus* isolates can survive and even grow at slow rates at supra-MIC concentrations of the fungicidal drug voriconazole.

## RESULTS

### Some *Aspergillus fumigatus* isolates show persistence to voriconazole

To determine whether *A. fumigatus* can display tolerance or persistence to voriconazole, we examined a collection of isolates consisting of 9 environmental, 10 clinical (gift from Prof Paul Dyer, collection PD-47-XX, here shortened as PD-XX), 5 common laboratory strains and one resistant control (RC) harbouring the TR34/L98H mutations in the *cyp51A* locus (Table S1 in https://doi.org/10.5281/zenodo.7021623), using disc diffusion assays. We evenly spread 4×10^4^ conidia of the isolates on RPMI agar plates, placed a 6 mm disc in the centre, containing 10 μL of voriconazole (0.8 mg/mL), and incubated them for 5 days. We observed variability in the sizes of the inhibition halos, reflecting differences in susceptibility of the isolates (Figs. 1A and S1). As expected, the RC strain did not show a proper inhibition halo (Figs. 1A and S1). In contrast, 15 of the 20 isolates showed a clear and well-defined inhibition zone. Interestingly, we found that 5 isolates were able to form colonies within the halo of inhibition, and 4 in 5 of these isolates (PD-9, PD-104, PD-254 and PD- 266, Fig. S1) formed only a few small colonies, which might be indicative that a few conidia are able to germinate and grow a little in the presence of a supra-MIC concentration of voriconazole (Fig. 1A and S1). The remaining isolate (PD-256) did not show any inhibition halo, suggesting that it is a resistant isolate.

**Figure 1.**
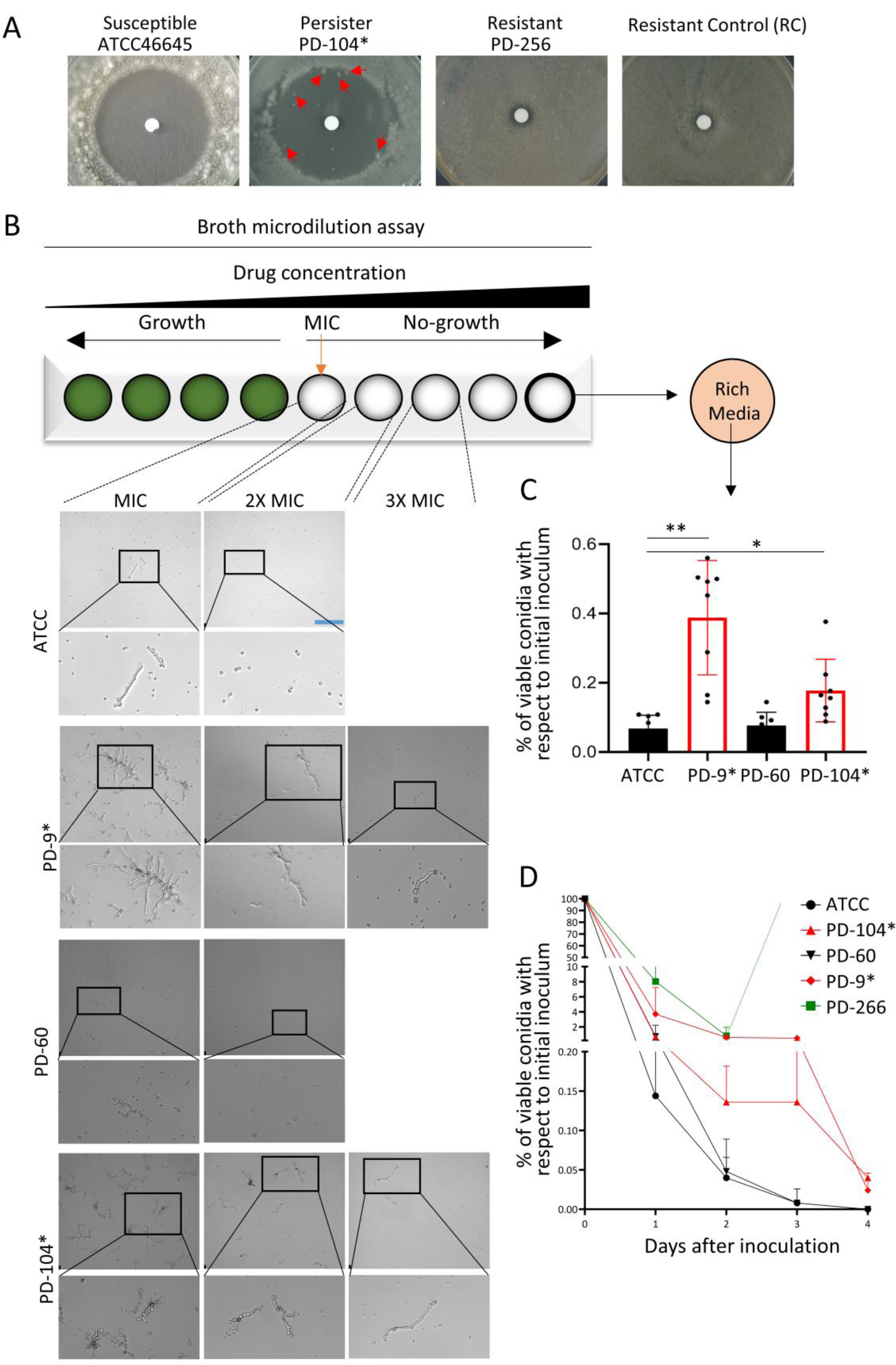
Certain *Aspergillus fumigatus* isolates display persistence to voriconazole. **A)** In disc-diffusion assays (10 µL of 0.8 mg/mL voriconazole), the susceptible isolate ATCC46645 never grew any colony in the inhibition halo, the resistant control (RC) isolate grew up to the edge of the disc, as did the strain PD-256. The persister isolate PD-104 was consistently able to grow a few small colonies. Plates were incubated for 5 days at 37°C. **B)** Inspection of the wells of a broth dilution assay under the microscope 72 hours after inoculation revealed that the non-persister isolates ATCC and PD-60 displayed only limited microscopic growth at the MIC concentration, and all conidia remained non-germinated at higher concentrations. In contrast, the persister strains PD-9 and PD-104 showed noticeable microscopic growth up to three fold (3x) the MIC concentration. Scale bar= 132.5 µm. **C)** Full content of the well containing the maximum concentration of voriconazole (8 µg/mL) was plated on rich media PDA plates and (CFUs) counted 48 hours after inoculation. Persister isolates grew significantly more CFUs than non-persister isolates (PD-9 VS ATC46645C *p*=0.002 and PD-104 VS ATCC46645 *p*=0.0331), demonstrating that these strains remain viable upon azole treatment for a longer period. Three independent experiments with three biological replicates were performed, the graph represents the means and SD, and data was analysed using the Brown-Forsythe and Welch ANOVA test with Dunnett’s multiple comparisons. **D)** A survival curve in the presence of 4 µg/mL of voriconazole revealed that, whilst non-persister strains lost viability very rapidly, the persister isolates had the characteristic bi-phasic reduction in viability, showing that a sub-population of the persister isolates remained viable for a longer period. Two independent experiments with biological duplicates and two technical replicates were performed; the graph represents the means and SD.

We then assessed the MICs of the original isolates and their derived colonies of the halo (CoHs). A complication of working with a filamentous fungus is that the CoHs need to be grown on a new plate in order to harvest spores (conidia) to be used for MIC determination. We therefore decided to assay two different conidia-harvesting conditions to distinguish the different phenotypes: re-growing the CoHs on solid RPMI in the absence or in the presence of a low concentration of voriconazole (0.12 µg/mL). It would be expected that for conidia grown in the presence of the drug, both resistance and heteroresistance are detected as an increment in the isolate’s MIC. In contrast, for conidia grown in the absence of the drug, the reversible increase in MIC that is characteristic of heteroresistance would be lost, whilst a stable genetic-based increment in MIC that defines resistance would be maintained. Finally, persistence should not cause a change of the isolate’s MIC independently of the presence of the drug in the conidia-harvesting medium. Using conidia obtained from both conditions, absence and presence of voriconazole, we found that the original isolate did not show an inhibition halo (PD-256, Fig. 1A and S1) had a very high MIC (>8 µg/mL), demonstrating that it is a resistant isolate (Table 2). CoHs formed by PD-254 and PD-266, which upon re-inoculation did not show an inhibition halo (Fig. S1), showed an increased MIC compared to the parental isolate (Table 2). The colony picked from PD- 254 showed increased MIC when re-grown on both media with and without voriconazole, suggesting that the CoH may have acquired a mutation that confers a stable resistance phenotype. In contrast, PD-266, which already had an elevated MIC, only showed an increased MIC when re-grown on medium containing voriconazole, suggesting that this strain may be heteroresistant. Finally, CoHs from isolates PD-9 and PD-104 showed the same MICs as their original isolates after a passage in the absence or presence of voriconazole (Table 2). This suggests that they are not resistant or heteroresistant derivatives of the original isolates. Indeed, repetition of the disc assay with the original isolates and with CoH re-grown in the presence of voriconazole, showed a similar level of colony appearance in the original strains and their derived CoHs for PD-9 and PD-104 (whereas it grew to the edge of the disc for PD-266, reflecting again a transient increase in its MIC) (Fig. S1). Although no formal measurement of growth rate was performed, the clear difference in size between single persister (PD- 9 and PD-104) and heteroresistant colonies (PD-266) after the same time of incubation (Fig. S1) indicate that the growth rate of persister colonies is reduced. Interestingly, all isolates showed a small increase (one dilution) in MIC when re-grown in the presence of drug (Table 2). This may be due to the development of conidia adapted to the growing environment in preparation for the subsequent germination, an effect recently described in *Aspergillus spp* [32]. Next, we tested if CoHs could also be detected using E-tests strips, which is a well-established method to measure the MIC and is thus more quantitative than disc diffusion assay [33]. In agreement with disc diffusion assays, CoHs developed inside the halo of inhibition created by E-tests on persister isolates (PD-9 and PD-104), but not on non-persister isolates (ATCC and PD-60) (Fig. S2A). Finally, to verify that the persistence phenotype is stable, we sequentially passaged CoHs isolated from PD-9 and PD-104 on PDA media (without voriconazole) and performed disc diffusion assays every two/three passages. The halo of the disc and the relative level of persistence were maintained for 10 passages (Fig. S2B), demonstrating that persistence is a stable phenotype.

**Table 2.** Voriconazole MICs of original isolates and their derived CoHs, pre-grown both in the absence and presence of a low (0.12 µg/mL) concentration of voriconazole

| Isolate | Voriconazole MIC ($\mu\text{g/mL}$) | | | | Classification of CoH |
| --- | --- | --- | --- | --- | --- |
|  | original | Original (grown on Drug) | CoH (- Drug) | CoH (grown on Drug) |  |
| ATCC | 1 | 2 |  |  | Susceptible |
| PD-9* | 0.5 | 1 | 0.5 | 1 | Persister |
| PD-104* | 1 | 2 | 1 | 2 | Persister |
| PD-254 | 2 | 2 | >8 | >8 | Resistant |
| PD-256 | >8 | >8 | >8 | >8 | Resistant |
| PD-266 | 4 | 8 | 4 | >8 | Heteroresistant |

In bacteria, persister and tolerant cells are often dormant and do not grow [19]. To investigate this in *A. fumigatus* in more detail, we followed a protocol to detect tolerance/persistence in bacteria in which the disc containing the drug, after a period of incubation, is switched, with another one containing fresh media [34]. The rationale is to detect surviving cells that have remained dormant and resume growth after the drug is withdrawn. After initial growth we substituted the disc with voriconazole for a new one containing *Aspergillus* Minimal Media (AMM). Interestingly, after the disc switch colonies were observed in the halos for several isolates (CEA10, PD-7, -8, -50, -154, -249, and - 264, Fig. S3), suggesting that these may be low level persister isolates. Nevertheless, three isolates, PD-9, -104 and -259, were able to form colonies before the disc switch, and of these three, only PD-9 and PD-104 grew even more colonies after disc switch (Fig. S3). Therefore, the isolates PD-9 and PD- 104 were able to survive and grow at slower rates in the presence of supra-MIC concentrations of voriconazole, a phenomenon that complies with the definition of persistence.

To further characterize persistence in *A. fumigatus*, we performed MIC assays with selected strains and checked microscopic growth of persister and non-persister isolates at supra-MIC concentrations. We found that 72 hours after inoculation the non-persister strains ATCC46645 and PD-60 displayed microscopic growth only at the MIC concentration, and no growth at all at higher concentrations (Figs. 1B and S4). In contrast, the persister isolates PD-9 and -104 showed noticeable microscopic growth at 2X, and even at 3X MIC germinated conidia could be detected (Fig. 1B and S4). While the level of growth was variable among independent experiments, the difference between persister and non-persister isolates was consistently observed (Fig. 1B and S4). In addition, we plated the entire content of the wells containing the highest concentrations of voriconazole (8 µg/mL) and found that 48 h after inoculation the non-persister strains were nearly all killed (only ∼0.07% of the conidia survived) whilst in the persister strains a significantly higher ratio of conidia survived (0.39% of PD-9 *p*=0.002 and 0.18% of PD-104 *p=*0.0331, Fig. 1C). Therefore, a subpopulation of the *A. fumigatus* persister isolates can survive for long periods in the presence of high concentrations of voriconazole and even grow at a slow rate, which seem to be inversely correlated with the concentration of drug.

In bacteria, a combination of MIC and speed of killing is the best approach to reveal and differentiate tolerance and persistence [18]. Therefore, we investigated the dynamics of cell death caused by voriconazole for persister and non-persister isolates. Conidia from two susceptible (ATCC46645 and PD-60), two persister (PD-9 and PD-104) and the putative heteroresistant (PD-266) strains were incubated for 4 days in liquid RPMI in the presence of 4 µg/mL of voriconazole and aliquots of the culture plated (after PBS washing to remove the drug) on PDA rich media every 24 hours (Fig. 1D). We found that for all strains the number of viable conidia dramatically declined after 24 hours of incubation, reflecting a strong fungicidal action of voriconazole, even against an isolate with high MIC. Interestingly, whilst the susceptible isolates were completely killed within 48-72 hours, a considerable number of conidia from the persister strains maintained viability for an extended period (∼0.748% for PD-9, ∼0.136% for PD-9 at 48 h and ∼0.52% for PD-9, ∼0.128% for PD-9 at 72 h, Fig. 1D), resulting in the characteristic biphasic killing curve that is the hallmark of persistence [19]. The isolate PD-266 showed visible growth from 48h, which agrees with its proposed heteroresistant phenotype, and saturated the plate, making impossible to count colony forming units (CFUs). To investigate the survival of the isolates at the single cell (conidial) level, we incubated the two persister and one susceptible strains in liquid RPMI in the presence of 32 µg/mL voriconazole for 48 hours, and replaced it with fresh drug free medium. We observed that during the first 24 hours of incubation after drug withdrawal there was no growth for any of the isolates, but subsequently the persister strains were able to resume growth (Videos S1 and S2 in https://doi.org/10.5281/zenodo.7021623) whilst the susceptible strain was not (Video S3 in https://doi.org/10.5281/zenodo.7021623). Hence, persister strains indeed can survive for extended periods in the presence of high concentrations of voriconazole and resume growth when the drug is withdrawn.

To check if the observed phenomena could be explained by mutations in the azole target genes, we sequenced the promoter and ORF of *cyp51A* and the ORF of *cyp51B* in the persister isolates PD-9 and PD-104 and the non-persister strain PD-60. In all strains we found that *cyp51A* was completely wild-type (there were no nucleotide changes) and *cyp51B* had only synonymous polymorphisms (Table S2 in https://doi.org/10.5281/zenodo.7021623), thus denying an implication of target enzyme mutations in the phenomenon of persistence. We also sequenced the sterol-sensing domain of *hmg1*, as it has been proposed that mutations in this gene may be a precursor step for the development of azole resistance [35, 36], and in bacteria persistence has been shown to correlate with the evolution of resistance [37, 38]. However, we found again that all strains harboured a completely wild-type sequence (not shown), suggesting that *hmg1* is not related with voriconazole persistence.

In conclusion, we have observed that certain *A. fumigatus* isolates can survive and grow at slow rates for extended periods of time in the presence of supra-MIC concentrations of voriconazole, therefore, these isolates display persistence to voriconazole.

### *Aspergillus fumigatus* persistence is maintained in the hyphal transition and seems to be medium dependent

*A. fumigatus* undergoes morphological changes during its development, shifting from resting conidia to swollen conidia (∼4-5 h), then to germlings (∼8 h) and finally forming hyphae (∼16 h) (Fig. S5A). As the cellular metabolic status is different at each developmental stage [39–41], we speculated that it might influence the capacity of the fungus to deploy persistence in the presence of voriconazole. Initially, to rule out that persistence could be explained by basal differences in germination or growth, we examined the germination and growth rates of two persister (PD-9 and PD-104) and two non-persister (ATCC and PD-60) isolates. We found that all isolates germinated at equivalent rate and ratio (Fig. S5B) and grew similarly on both solid (Fig S5C) and liquid (Fig. S5D) RPMI media. Once excluded differences in germination or growth, we tested the influence of the morphological stage on persistence by incubating the RPMI plates inoculated with the fungus for 8 or 16 hours before adding the drug to the disc. We found that the persister strains PD-9 and PD-104 were also able to form colonies in the halo when drug was added 8 hours after the beginning of incubation (Fig. 2A), suggesting that persistence is not determined by the developmental stage. We could not draw definitive conclusions for the 16 hours grown hyphae, as the background growth was too dense to undoubtedly differentiate specific persister colonies (Fig. 2A). Nevertheless, we could clearly detect a growing colony in the voriconazole halo for the strain PD-104 (Fig. 2A), proving that at least this strain can display persister growth when short hyphae are challenged with the drug. Interestingly, it has previously been reported that morphological status alters *A. fumigatus* susceptibility profiles against various drugs including voriconazole [42], which supports the notion that persistence is a different phenomenon, triggered by distinct mechanisms, and independent of MIC.

**Figure 2.**
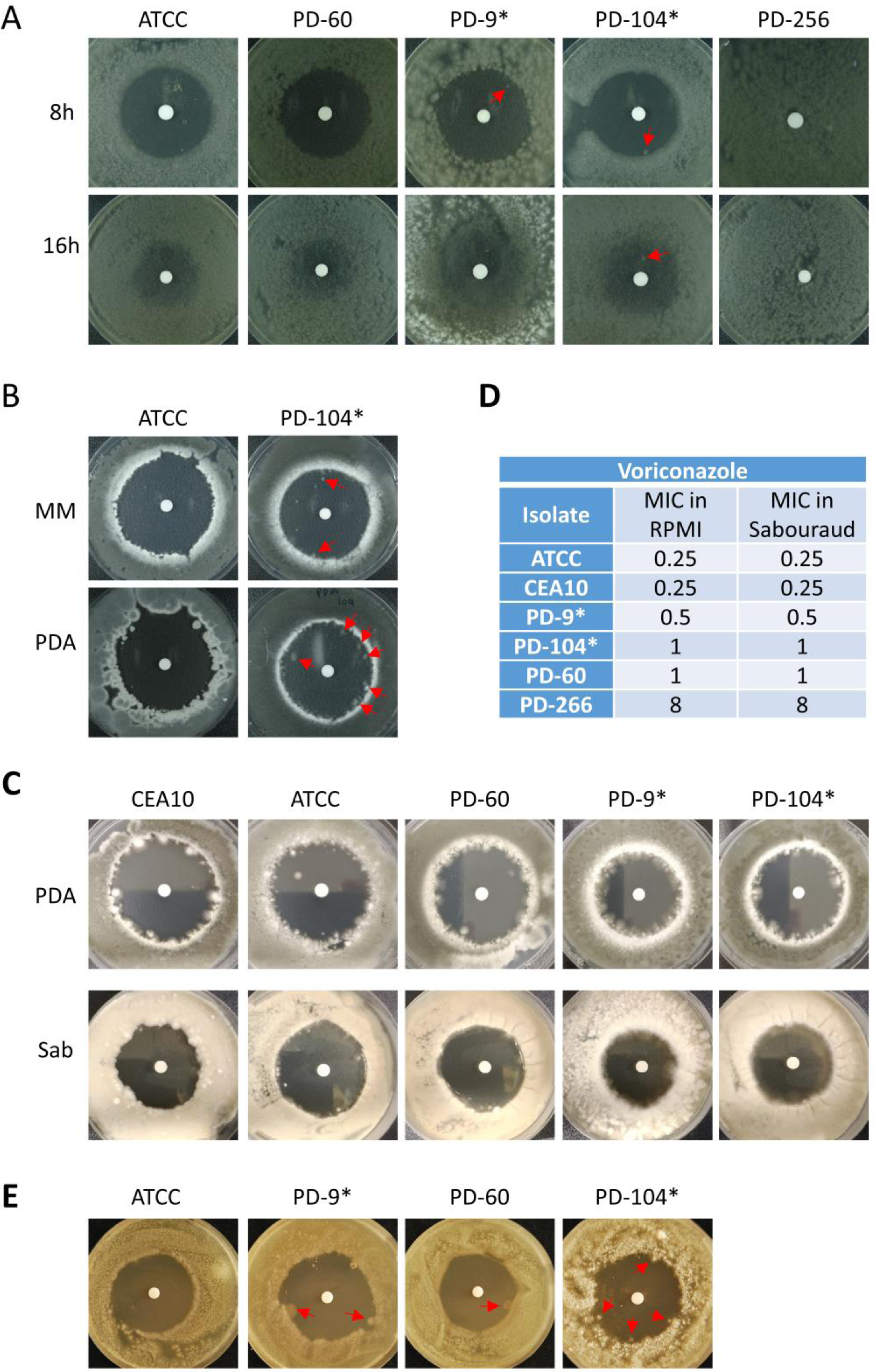
*Aspergillus fumigatus* persistence does not depend on the morphological stage, but seems to be determined by the growth media. **A)** Conidia of the different isolates were inoculated and incubated for 8 or 16 hours before the disc containing voriconazole (10 µL of 0.8 mg/mL) was added to the RPMI plate. At 8 hours, when conidia have germinated, the persister isolates PD-9 and PD-104 were still able to grow small colonies in the inhibition halo, whilst the non-persister strains were not. At 16 hours, when conidia have already formed hyphae, the background growth made impossible to distinguish colonies in PD-9, but a clear one could be detected in PD-104. **B)** On *Aspergillus* minimal medium (MM), the persister isolate PD- 104 was able to form colonies in the halo, but the non-persister strain ATCC was not. **C)** On rich media PDA and Sabouraud (Sab) all isolates were able to form conidia in the halo. **D)** This happened even the MIC of all isolates was the same in commonly assayed RPMI and the rich media Sab. **E)** 10-week old spores displayed a similar persister phenotype as freshly isolated spores. Often a single colony could be detected in the halo of inhibition on the non-persister PD-60, suggesting that aged spores might have slightly more persistence capacity.

To determine if persistence is influenced by the growth medium, we performed a voriconazole disc diffusion assay with the PD-104 isolate and the wild-type control ATCC46645 on PDA rich media or AMM. The PD-104 strain showed persister growth on both media, and it seemed to be able to form a higher number of COHs on PDA rich media (Fig. 2B). The ATCC46645 wild-type strain did not show persistence on AMM, but surprisingly it did form colonies in the halo on PDA media (Fig. 2B). To investigate this in more detail we repeated the disc assay using two different rich media, PDA and Sabouraud, with more strains, CEA10, PD-9, and PD-60 (Fig. 2C) and found that all strains were able to form colonies in the halos. The MICs of the isolates, measured by broth dilution assay, were equal on Sabouraud and RPMI (Fig. 2D), indicating that the effect of the drug is not reduced in rich medium. Therefore, the ability of *A. fumigatus* to display persistence seems to be medium dependent, suggesting that a rich nutrient environment favours survival at supra-MIC concentrations. This is supported by a recent study that showed that a rich metabolic environment can promote azole tolerance in *Saccharomyces cerevisiae* [43]. Nevertheless, it should also be considered that voriconazole diffusion may be affected in solid rich media, which could be confounding this observation.

Finally, we also considered that persistence might be affected by the age of the spores (all experiments are performed with freshly harvested spores). To check this, we assayed disc diffusion assays using 10-weeks old spores. We did not observe remarkable differences with respect to fresh spores, although in most of the experiments we could see one colony in the halo of the non-persister PD-60 (Fig. 2E), suggesting that aged spores might have a slightly higher persistence potential, at least for some isolates.

### *Aspergillus fumigatus* persistence to voriconazole is independent of stress and cannot be inhibited with adjuvant or antifungal drug

Changing environmental conditions activate signalling cascades that trigger transcriptional adaptation and cell wall alterations [44–47]. Therefore, we reasoned that the capacity of certain isolates to survive and grow in supra-MIC concentrations of voriconazole might be influenced by environmental stressors. Indeed, in *C. albicans* mutants and inhibitors of stress response pathways eliminate tolerance [48]. To investigate this possibility, we analysed voriconazole persistence of PD-104 in the presence of hypoxic (1% O_2_), oxidative (0.01% H_2_O_2_), osmotic (150 mM NaCl), membrane (0.05% SDS) and cell wall (10 µg/mL CalcoFluor White) stress. Surprisingly, in contrast to *C. albicans* [21], we found that most environmental conditions did not influence persistence in *A. fumigatus* (Fig. 3A). This suggests that the underlying mechanism(s) of persistence in *A. fumigatus* are likely different from the previously proposed mechanisms of tolerance in *C. albicans*. The only condition that influenced *A. fumigatus* persistence was hypoxia, which could prevent growth in the halo (Fig. 3A). As we had observed above that persistence was influenced by the growth medium, we wondered if hypoxia could prevent persistence also on rich medium. As shown in Figs. 2B and 2C, all isolates were able to grow colonies in the halo when grown on the rich medium YAG on normoxia (Fig. 3B). However, persistence was eliminated under hypoxia for all strains, except for PD-104, for which it was reduced but apparently not completely eradicated. Therefore, it seems that hypoxia reduces persistence, but the nutritional composition of the medium also influences its impact on the phenomenon.

**Figure 3.**
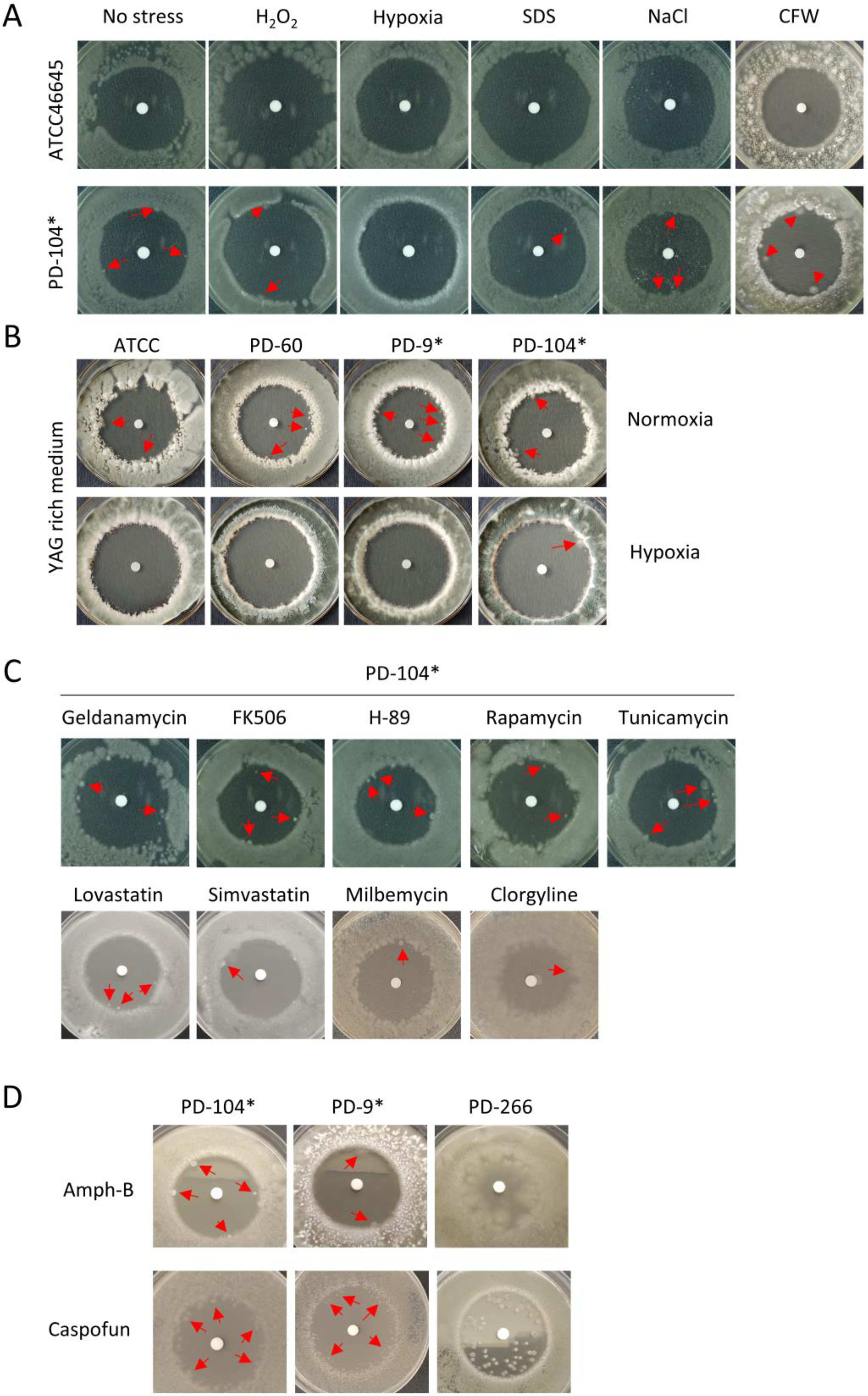
*Aspergillus fumigatus* persistence is not influenced by stress, with the exception of hypoxia, and cannot be eliminated with adjuvant or combinatorial treatments. **A)** Hypoxia was the only stress that could prevent persistence, whilst neither oxidative (H_2_O_2_), cell wall (SDS or CFW) or osmotic (NaCl) stress influenced persistence. **B)** Hypoxia completely eradicated persistence also on the rich medium YAG for ATCC, PD-60 and PD-9, but only reduced persistence for the PD-104. **C)** The use of adjuvant drugs could not prevent persistence. **D)** Combinatorial treatment with amphotericin-B and caspofungin did not prevent persistence. All plates were incubated with 10 µL of 0.8 mg/mL voriconazole added to the disc and the specific condition (stress, adjuvant or combinatorial drug) as described in the text. Plates were incubated for 5 days at 37**°**C. All plates and conditions were repeated in at least two independent experiments.

In *C. albicans*, fluconazole tolerance (but not resistance) can be prevented with the use of adjuvant drugs that block general stress signalling pathways [21]. To further investigate if the underlying mechanism of persistence may be different in *A. fumigatus*, we tested the effect of various drug adjuvants that were previously shown to eliminate tolerance in *C. albicans* [21]: geldanamycin (0.8 µg/mL), an inhibitor of heat shock protein (Hsp90) [49, 50], FK506 (4 ng/mL), an inhibitor of calcineurin [51], H-89 (4 µg/mL), an inhibitor of the cAMP-dependent protein kinase (PKA) [52], rapamycin (6.25 µg/mL), an inhibitor of the mammalian target of rapamycin (mTOR) [53], and tunicamycin (10 µg/mL), an inducer of the unfolded protein response pathway [54]. In contrast to *C. albicans*, the use of adjuvant drugs did not prevent persistence of *A. fumigatus* isolates (Fig. 3C). We also tested lovastatin (8 µg/mL) and simvastatin (2 µg/mL), as statins inhibit 3-hydroxy-3-methylglutaryl-coenzyme A (HmgA) [55], an enzyme in the same metabolic pathway as the target of azoles, and these two were previously shown to have antifungal activity against *Aspergillus spp* [56]. However, statins were also not able to prevent persister growth (Fig. 3B). Finally, as efflux of antifungals has been proposed to play a role in *C. albicans* tolerance [48], we tested if the efflux inhibitors milbemycin A oxim (8 µg/mL) [57] or clorgyline (63.5 μg/mL) [58] can prevent *A. fumigatus* persistence. These compounds seemed to be able to diminish the persister capacity of the strains, as only one CoH per plate could be detected, and these colonies were exactly at the edge of the halo (Fig. 3C). In conclusion, adjuvant drugs cannot prevent *A. fumigatus* persistence to voriconazole, and efflux inhibitors may affect this process and deserve further study.

Next, we evaluated if persistence can be eradicated using combinatorial treatment with the other classes of antifungal drugs in clinical use. However, neither amphotericin-B (1 µg/mL) nor caspofungin (0.5 µg/mL) could prevent persistence (Fig. 3D). Interestingly, these antifungal drugs could also not impede growth in the halo of the presumed heteroresistant strain PD-266 (Fig. 3D). Therefore, it seems that combinatorial treatment with other antifungals cannot prevent persistence in *A. fumigatus*.

### The phenomenon of persistence can be observed with other azole drugs

To check if *A. fumigatus* can display persistence in the presence of other azoles, we firstly employed the disc diffusion assay adding 10 µL of a 3.2 mg/mL itraconazole solution. We found that the isolates PD-104 and PD-266 were able to form colonies in the halo (Fig. S6). Upon re-inoculation of a CoH (re- grown in itraconazole containing media), the isolate PD-104 showed an inhibition halo of the same size and displayed similar level of CoH appearance, whereas the PD-266 isolate was able to grow on the whole plate and did not show any inhibition halo. Therefore, this suggests that, as observed with voriconazole, the isolate PD-104 is persistent and the isolate PD-266 is heteroresistant to itraconazole. We then performed a broth dilution assay with our four well-characterised isolates (non-persisters ATCC46645 and PD-60 and persisters PD-9 and PD-104) to calculate the MIC for itraconazole and isavuconazole, and looked under the microscope at supra-MIC concentrations (Fig. S7A and S7B). We found that the isolates PD-9 and PD-104 showed slight growth at 2X MIC, whilst ATCC46645 and PD- 60 did not (Fig. S7B). Moreover, we inoculated the entire content of wells containing the highest concentration of the drugs (8 µg/mL) on PDA plates, and found that a detectable number of conidia from the isolates PD-9 and PD-104 remained viable after 48 hours of incubation in the presence of the azoles (Fig. S7C). Hence, even if more experiments need to be done for a detailed characterization of persistence to these azoles, these results suggest that the same isolates that are persisters to voriconazole could also display persistence to itraconazole and isavuconazole.

### The transcriptome of persister growth suggest that Galactosaminogalactan and high expression of sterol biosynthetic genes may be relevant to establish persistence

To better understand the heterogeneous nature of persistence, we compared the transcriptome of PD-104 grown under conditions favouring persistence (above MIC), sub-MIC and in the absence of drug. We inoculated the spores on top of a nylon membrane placed on an RPMI solid plate. We put the disc with voriconazole (0.8 mg/mL) on the membrane and incubated for 5 days at 37°C. Standard halos of inhibition formed on the membrane, and persister colonies appeared for the PD-104 strain, but not for the A1160 strain (Fig. S8A). We harvested mycelium (avoiding conidia as much as possible) from plates without a voriconazole disc (No Drug), from the area equidistant from the border of the plate and the inhibition halo (Low Drug) and the colonies in the halo (Persistence) (Fig. 4A). We had to combine persister colonies from 20 plates per replicate in order to obtain sufficient material for RNA extraction. Additionally, we harvested mycelia from A1160 with No and Low Drug under the same conditions. We performed RNA-seq of two biological replicates/condition and compared the transcriptomes of the following conditions (i) A1160 Low Drug VS No Drug, (ii) PD-104 Low Drug VS No Drug, (iii) PD-104 Persistence VS No drug and (iv) PD-104 Persistence VS Low Drug. We considered as differentially expressed genes (DEG) those with a log2FoldChange >1 or <-1 and a false discovery rate (FDR) <0.05.

**Figure 4.**
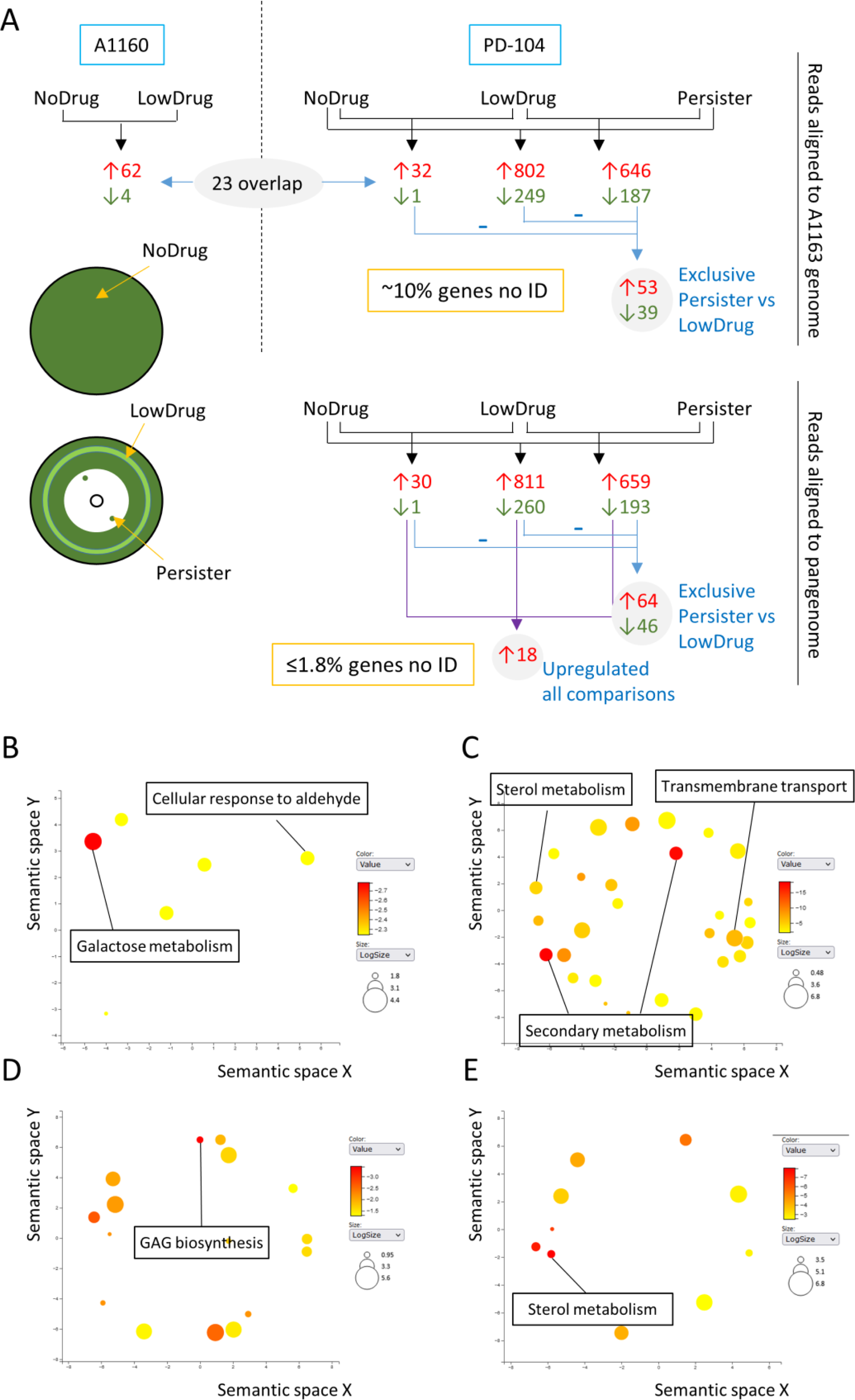
RNA-seq analysis reveals potential causes of persistence. **A)** Schematic representation of the conditions assayed, the comparisons made and the number of genes identified, as detailed in the text. In addition, the sites of sampling for the no drug (plate without voriconazole disc), low drug (light green circle at mid-distance of the inhibition halo) and persistence (colonies inside the inhibition halos) conditions are shown. **B-E** The most significantly upregulated GO biological processes are shown using the REVINGO tool for GO data visualization [67]. The full list of upregulated GO terms can be found in Tables S2-S5 and Table 3. **B**) PD-104 Persister only genes (aligned to the A1163 genome), **C)** all genes upregulated in Persistence (aligned to the pangenome), **D)** PD-104 Persister only genes, (aligned to the pangenome) and **E)** genes that are most upregulated in persistence compared to normal response to the drug.

In A1160, we detected 62 genes significantly upregulated in the presence of low voriconazole concentrations and only 4 that were downregulated (Fig. 4A and Dataset S1 in https://doi.org/10.5281/zenodo.7021623). Such a low number of DEGs may be due to the relatively low concentration of voriconazole in the harvested area. Nevertheless, among the upregulated genes we found 10 related to sterol biosynthesis (Dataset S1 in https://doi.org/10.5281/zenodo.7021623), including *cyp51A* and *cyp51B*, the expression of which have been reported to increase upon azole challenge [59, 60]. Surprisingly, only 21 of the 62 upregulated genes overlapped with our previously published dataset, in which 1492 genes were upregulated upon challenge of A1160 with itraconazole [61]. This striking difference may be due to the different drug employed, the different concentrations assayed and/or the use of solid medium. Interestingly, these 41 DEGs that are specific to this analysis (which can be found in Dataset S1 in https://doi.org/10.5281/zenodo.7021623) contain *cyp51A*, *cyp51B* and various other genes of the sterol pathway.

For PD-104, the Low Drug VS No Drug comparison retrieved 32 genes upregulated and only 1 downregulated (Fig. 5 and Dataset S2 in https://doi.org/10.5281/zenodo.7021623). Within the upregulated genes we again found 8 genes related with sterol biosynthesis, including *cyp51A*. In addition, 9 genes did not overlap with the comparison of A1160 (Fig. 4A and Dataset S2 in https://doi.org/10.5281/zenodo.7021623). These differences with A1160 may point to potential strain differences in response to voriconazole. In the Persister VS No Drug comparison there were 802 upregulated and 249 downregulated genes and in the Persister VS Low Drug 646 upregulated and 187 downregulated genes (Fig. 4A and Dataset S2 in https://doi.org/10.5281/zenodo.7021623). When comparing those three analyses, only 53 genes were exclusively upregulated and 39 downregulated in Persister VS Low Drug (detected using BioVenn [62]), suggesting that they may be important to establish the persister phenotype and not only as a response to the drug (Fig. 4A and Dataset S2 in https://doi.org/10.5281/zenodo.7021623). We performed Gene Ontology (GO) enrichment analyses (on FungiDataBase [63]) for these 53 DEGs, which retrieved a number of biological processes and molecular functions for up- and downregulated genes. It is important to note that the FDR did not reach significance for any of them, which is possibly due to the low number of genes included in the analysis. However, we believe this analysis provided interesting clues and can help to direct future research. For the up-regulated genes, the strongest enrichment was for the biological processes galactose and aldehyde metabolisms (Fig. 4B) and the molecular function oxidoreductase activity (Table S3 in https://doi.org/10.5281/zenodo.7021623). Interestingly, galactose metabolism was found to be the most upregulated function in *Staphylococcus aureus* persisters, although the reason has not been elucidated yet [64]. For the downregulated genes, there was an enrichment of biological processes related with development and cell cycle (Table S3 in https://doi.org/10.5281/zenodo.7021623), possibly reflecting the reduced growth rate in this condition.

**Figure 5.**
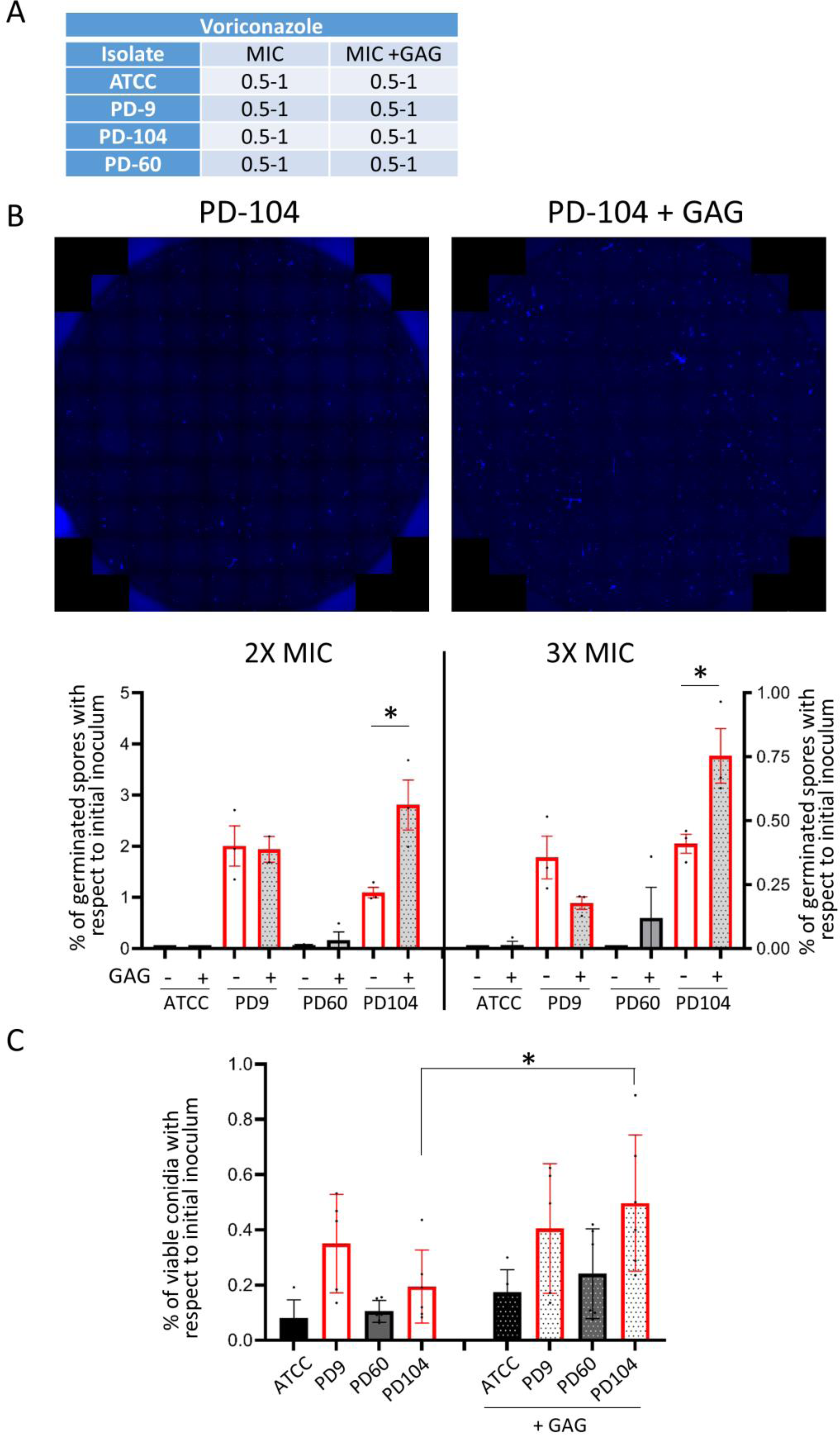
External addition of GAG potentiates persistence in the isolate PD-104. **A)** Addition of GAG does not change the MIC of the isolates. **B)** Representative merges of 87 photos to cover the whole well of broth dilution assays. The fungal material was stained with calcofluor white. Representative wells of PD-104 with and without GAG at 2X the MIC are shown **C)** Counting of germinated conidia/short hyphae in the entire wells of 2X (left) and 3X (right) MIC of broth dilution assays. External addition of GAG significantly increased the number of germlings in PD-104 from 1.09% to 2.80%, *p=*0.0267 at 2X MIC and from 0.41% to 0.75% *p*=0.038, two-tailed unpaired *t*-test. Two independent experiments with three technical replicates were performed. The graphs represent the means and SD. **D)** Plating of the entire content of wells containing 4 µg/mL of voriconazole shows that external addition of GAG significantly increased the number of viable PD-104 cells that can be recovered 48 hours after inoculation (*p*=0.0437, one way ANOVA with Tukey’s multiple comparisons). Two independent experiments with three biological replicates were performed. The graph represents the means and SD.

Interestingly, a significant number of the detected differentially expressed mRNAs in PD-104 were not assigned to any gene of the A1163 genome reference used for the mapping of the reads. In detail, 80 of the upregulated (including 4 of the 20 most upregulated) and 29 of the downregulated for the Persister VS no Drug comparison and 55 upregulated (including 5 of the 10 most upregulated) and 14 downregulated for the Persister VS no Low comparison seem to have no orthologue in A1163 (Dataset S2 in https://doi.org/10.5281/zenodo.7021623). This points to the requirement of a high number (∼10% of the DEG) of strain specific genes to facilitate for persister growth. Therefore, we re-annealed the sequenced mRNAs from the PD-104 isolate to the newly generated *A. fumigatus* pangenome [65] and performed the comparisons again (Dataset S3 in https://doi.org/10.5281/zenodo.7021623). In this analysis, the Persister VS no Drug comparison retrieved 811 genes up- and 260 downregulated, and the Persister VS Low Drug comparison 659 up- and 193 downregulated (Fig. 4A and Dataset S3 in https://doi.org/10.5281/zenodo.7021623). The ratio of identified genes greatly improved, as only ≤1.8% of the DEGs could not be assigned to an ID. Yet, the most upregulated gene in the Persister VS Low Drug comparison was an unidentified gene and many of the newly identified genes encode uncharacterised hypothetical proteins, which reflects our ignorance of *A. fumigatus* biology and thus how difficult it is unravelling underlying mechanisms in this pathogen. GO enrichment analysis of all DEGs showed that secondary metabolism, ergosterol metabolism and transport were key upregulated biological processes (Fig. 4C) and oxidoreductase, catalysis and transmembrane transport important molecular functions (Table S4 in https://doi.org/10.5281/zenodo.7021623), all of which suggests an active metabolic response in persister conditions. As observed above (Table S3 in https://doi.org/10.5281/zenodo.7021623), downregulated biological processes reflected a downregulation of developmental processes (Table S4 in https://doi.org/10.5281/zenodo.7021623). Comparison of the DEGs present in all three comparisons revealed that 64 genes were exclusively up- and 46 downregulated in Persister VS Low Drug (detected using BioVenn [62]), suggesting that they may be specifically important for persistence (Fig. 4A and Dataset S3 in https://doi.org/10.5281/zenodo.7021623). GO enrichment analysis of those genes revealed galactosaminogalactan (GAG) biosynthesis as the most significant upregulated biological process (Fig. 4D and Table S5 in https://doi.org/10.5281/zenodo.7021623), providing a plausible explanation for the upregulation of galactose metabolism observed before, and suggesting that this exopolysaccharide may be relevant for persistence in *A. fumigatus*. Next, we performed a protein functional association analysis using the STRING database and platform [66] to investigate if these 64 upregulated genes are functionally correlated. Sixty-one of these genes could be matched to proteins in the database, showing a significant (*p*=1.84e-08) interaction network, including a node of 17 proteins related with metabolism (Fig. S8B and Table S6 in https://doi.org/10.5281/zenodo.7021623). This suggests that a distinct metabolic response occurs during persister growth.

Bacterial persistence has been proposed to be a sub-population event due to stochastic high expression of relevant genes [68, 69]. Accordingly, we reasoned that those genes that are upregulated in Low Drug VS No Drug and Persister VS No Drug comparisons, but also in Persister VS Low Drug might reveal those genes that are important to adapt to presence of the drug, but also that can create persistence with higher levels of expression. We identified 18 genes that appeared as upregulated in all three comparisons (Fig. 4A and Table 3), indicating that they have higher levels of expression in persistence VS normal response to the drug. GO enrichment analysis revealed sterol metabolism as the most significantly upregulated biological process (Fig. 4E and Table S7 in https://doi.org/10.5281/zenodo.7021623). Indeed, those few genes had a very significant enrichment in the KEGG pathway “steroid biosynthesis” (Table 3), suggesting that high expression of genes in the sterol biosynthetic route (including *cyp51A*) may enable the sub-population of persisters to survive and grow in supra-MIC concentrations. Additionally, these highly expressed genes were enriched in KEGG pathways related with cytochrome P450 dependent drug metabolism (Table 3), which may indicate that detoxification of azoles is also important for persistence. Finally, expression of the *cdr1B* transporter (AFUA_1G14330), known to be associated with azole resistance [70], was also detected to be higher in persistence (Table 3). This suggests that a higher capacity to efflux azoles may also be important for the persister phenotype. Finally, we performed a protein functional association analysis with these 18 genes using the STRING database and platform [66] to search for functional correlations. All 18 genes were matched to proteins and a significant interaction (*p*<1.0e-16) was found, involving a highly interactive nodule of 6 proteins related with steroid biosynthesis (Fig. S8C and Table S8 in https://doi.org/10.5281/zenodo.7021623), suggesting again that a high production of ergosterol is important for persistence.

**Table 3.** GO biological processes enrichment and KEGG pathway enrichment for the 18 genes that showed higher expression in Persistence than in NoDrug and LowDrug.

| Gene ID | Product description |
| --- | --- |
| AFUA_1G03200 | Putative major facilitator superfamily (MFS) transporter |
| AFUA_4G06890 | 14-α sterol demethylase Cyp51A |
| AFUA_2G00320 | putative sterol delta 5,6-desaturase |
| AFUA_3G00810 | Putative cholestenol delta-isomerase |
| AFUA_1G03150 | C-14 sterol reductase, putative |
| AFUA_8G02440 | C-4 methyl sterol oxidase, putative |
| AFUA_6G14140 | Has domain(s) with predicted role in response to stress and integral component of membrane localization |
| AFUA_3G00150 | Has domain(s) with predicted oxidoreductase activity and role in metabolic process |
| AFUA_4G01440 | Predicted glutathione S transferase |
| AFUA_4G04820 | C-4 methyl sterol oxidase Erg25, putative |
| AFUA_5G00840 | Ortholog of *A. nidulans* FGSC A4 : AN5639, AN2587, AN9444, AN7395, |
| AFUA_2G15130 | ABC multidrug transporter A-2, putative |
| AFUA_5G03290 | Ortholog of *A. nidulans* FGSC A4 : AN8197, A |
| AFUA_3G02520 | Has domain(s) with predicted role in transmembrane transport and integral component of membrane localization |
| AFUA_7G04740 | Ortholog of *A. fumigatus* Af293 : Afu1g01960, Afu3g01040, Afu3g03315, Afu7g01960, Afu8g05750, Afu5g00135, Afu7g06526 |
| AFUA_2G01890 | Ortholog(s) have 2-octoprenyl-3-methyl-6-methoxy-1,4-benzoquinone hydroxylase activity, |
| AFUA_1G14330 | Azole transporter |
| AFUA_8G00710 | Has domain(s) with predicted role in defense response, negative regulation of growth |

| KEGG Pathway |  |  |  |
| --- | --- | --- | --- |
| ID | Name | P-value | Benjamini |
| ec00100__PK__KEGG | Steroid biosynthesis | 2,31E-08 | 1,18E-06 |
| ec00980__PK__KEGG | Metabolism of xenobiotics by cytochrome P450 | 0.002873 | 0.07326 |
| c00982__PK__KEGG | Drug metabolism - cytochrome P450 | 0.004423 | 0.07520 |

### Galactosaminogalactan potentiates persistence of PD-104

Our transcriptome analysis detected a significantly higher expression of two GAG biosynthetic genes (*sph3*, AFUA_3G07900 and *uge3*, AFUA_3G07910) in persister growth (Dataset S3 in https://doi.org/10.5281/zenodo.7021623). Manual inspection of the GAG biosynthetic genes [71, 72] revealed that another gene (*agd3*, AFUA_3G07870) might also be upregulated (FC= 1.01), but although the *p*-value was significant (*p*= 0.0179), the FDR rate was above threshold (FDR=0.1167) (Dataset S3 in https://doi.org/10.5281/zenodo.7021623). This prompted us to investigate if a higher amount of GAG could create and/or potentiate persistence in *A. fumigatus*. To this aim, we performed broth dilution assays in which we inoculated each isolate (ATCC, PD-9, PD-60 and PD-104) in two different lines and added purified GAG at a concentration of 100 µg/mL to one of them. Comparison of the MIC with and without GAG demonstrated that this polysaccharide did not affect the resistance profile of the isolates (Fig. 5A). After reading the MIC at 48 hours, we added 10 µg/mL of the dye calcofluor white (CFW) to the wells and imaged the entire well (87 photos using a 20X objective) at the blue emission channel. Images were merged (Fig. 5B) and the number of germinated conidia/short hyphae in the whole well calculated as explained in materials and methods (Fig. 5B). By these means we could calculate that ∼2% of the PD-9 spore population and ∼1% of the PD-104 conidia were able to germinate at 2X MIC, and ∼0.35% for both isolates in 3X MIC. Interestingly, addition of GAG enhanced the number of persisters in PD-104 (from 1.09% to 2.80%, *p=*0.0267 at 2X MIC and from 0.41% to 0.75% *p*=0.038) but not in PD-9 (Fig. 5B). Next, we inoculated the four isolates in duplicated lines of 96-well plates containing a high concentration of voriconazole (4 µg/mL) in all wells, and added GAG to one line of each isolate. The entire contents of wells were plated 48 hours after inoculation on PDA rich plates to count the number of viable CFUs. As observed before (Fig. 1C), we found that the persister isolates (PD-9 and PD-104) maintained viability of a significant number of conidia for an extended period of time (Fig. 5C). Interestingly, GAG addition seemed to slightly increase the number of grown CFUs for all isolates, although this increment was only significant for the PD-104 isolate (PD- 104 VS PD-104+GAG *p*=0.0437).

These results suggest that a high level of GAG can potentiate persistence in some, but apparently not all, isolates. Future investigations will aim to understand the underlying mechanism and to determine why GAG addition is isolate-specific.

### Persistence can be detected in diverse *A. fumigatus* collections of isolates

To corroborate that certain *A. fumigatus* isolates display persistence to voriconazole, we decided to screen two independent collection of isolates. Initially, we tested an environmental library of isolates collected in the area of Manchester, UK [73]. We screened all 157 isolates for growth under the microscope after 72 hours incubation in the presence of a high concentration of voriconazole (8 µg/mL). We found that 34 isolates were able to show a limited degree of growth, suggesting that they might be persisters (Table S9 in https://doi.org/10.5281/zenodo.7021623). Next, we employed disc diffusion assays with 0.8 mg/mL voriconazole and observed that 15 out of the 34 isolates were able to form colonies in the halo (Table S9 in https://doi.org/10.5281/zenodo.7021623). Broth dilution assay with the original isolates and the CoHs revealed that 10 of those isolates did not have increased MICs, indicating that they were persister strains (Table S9 in https://doi.org/10.5281/zenodo.7021623). Therefore, at least ∼6% of this collection was persister to voriconazole. We further evaluated a collection of 17 azole-susceptible clinical isolates obtained from TAU medical centre (Table S9 in https://doi.org/10.5281/zenodo.7021623), which were prior shown not to have any mutation in the *cyp51A* gene promoter or ORF, using the disc diffusion assay. We found that 6 out of 17 (35%) isolates were able to grow small colonies in the inhibition halo (Fig. S9A). Colonies picked from within the halo formed upon re-inoculation similar size halos, indicating that the MIC had not increased, and grew a similar number of CoHs, suggesting that the isolates are persisters.

Therefore, evaluation of independent collection of isolates seem to always retrieve a significant number of persister strains, demonstrating that this is quite a common phenomenon.

### Persistence is not a lineage specific trait

Recent studies suggest that the tandem repeats (TRs) in the promoter region of *cyp51A*, which cause high levels of antifungal resistance, possibly have evolved in the environment, due to the use of demethylase inhibitors (azoles) in the fields [74]. Interestingly, the isolates with the TR_34_/L98H polymorphism have been found to be closely related [65, 74], which suggest that this mechanisms has evolved in a distinct lineage of the species. To investigate if the phenomenon of persistence could also be lineage specific, we sequenced the genome of 23 of our isolates, 12 persisters and 11 non-persisters (Table S9 in https://doi.org/10.5281/zenodo.7021623) and integrated their genome into a recently published phylogenetic analysis [65]. We found that both persister and non-persister isolates scattered through the entire tree, demonstrating that persistence is not a trait of a specific lineage of *A. fumigatus* (Fig. 6). In addition, it is noteworthy that some persister and non-persister isolates seem to be genetically very similar, as they are close in the tree (JA12-JA14-JA7-JA8 and also JA1-JA2, Fig. 6). This suggests that the genetic features that enable persistence are modest and/or that non-genomic features, as the transcriptional response, are important for isolate specificity.

**Figure 6.**
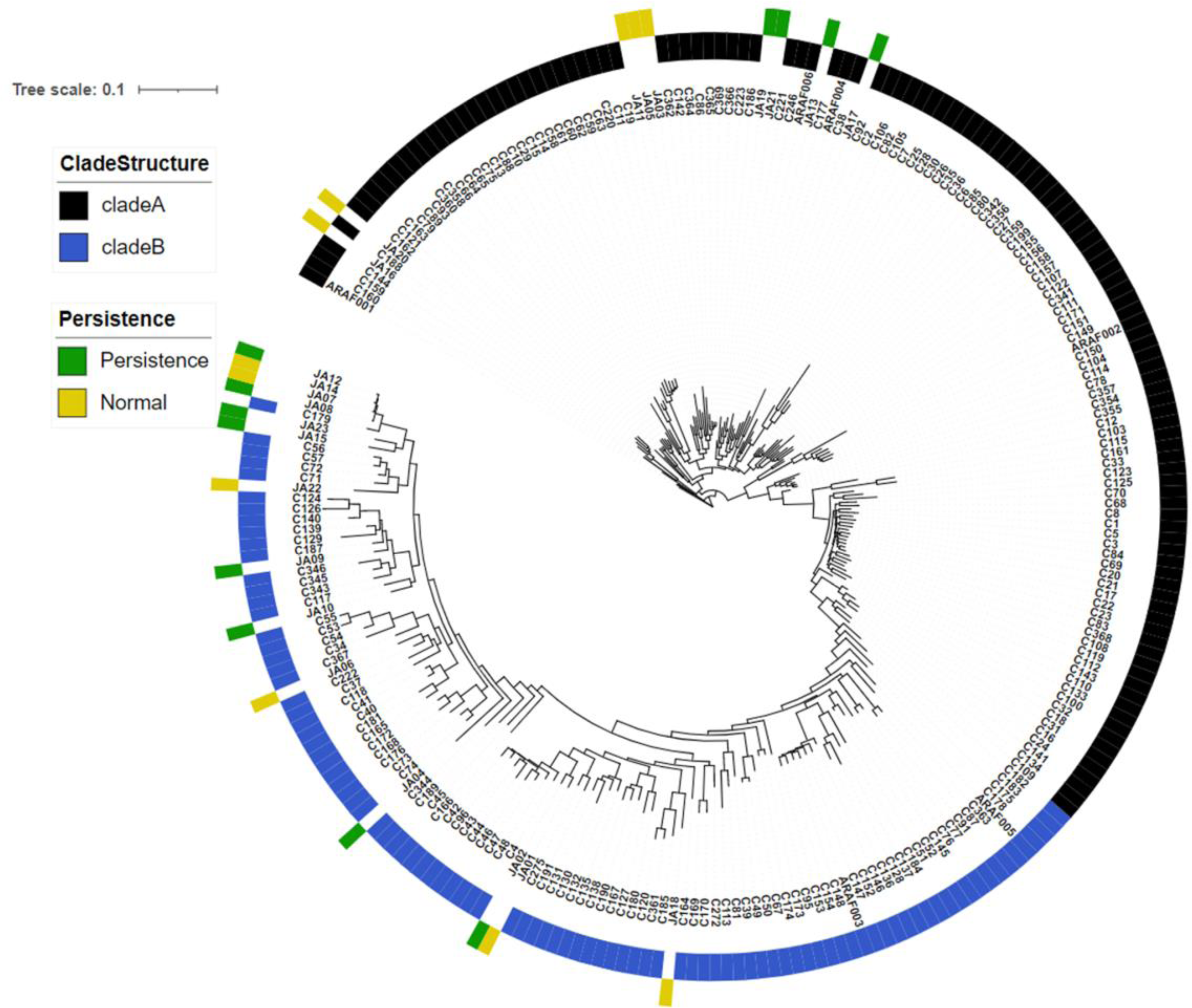
Persister isolates do not form a distinct lineage. The genomes of newly sequenced persister and non-persister isolates, as characterised in this study (Table S9 in https://doi.org/10.5281/zenodo.7021623), distributed scattered when included in the previously generated phylogenetic tree [65]. Therefore, persistence is not a feature that has evolved in a particular lineage.

### In a *Galleria mellonella* infection model, a voriconazole treatment seems to be less efficient against persister isolates in some larvae

To evaluate if persistence may be relevant during antifungal treatment, we employed the *Galleria mellonella* mini-host model of infection. This model has been successfully used to investigate the efficiency of azole treatment against *A. fumigatus* [75, 76]. In addition, a recent study has characterised the pharmacokinetics of voriconazole in infected larvae, which allowed us selecting the optimal dose to reach a high concentration of the drug in the haemolymph (above the MICs of the isolates), but that is nearly completely removed in 24 hours [77].

Initially, we performed a survival experiment with all four strains to investigate if they could have different virulence potential. We infected larvae with 10^4^ or 5×10^4^ conidia of non-persister (ATCC46645 or PD-60) or persister (PD-9 or PD-104) isolates and followed mortality for 10 days. All the isolates killed larvae at a similar rate (Fig. S9B), demonstrating that they are similarly virulent. Therefore, we aimed to use this model to investigate if persistence could potentially cause treatment failure. We reasoned that a voriconazole treatment should very efficiently eradicate susceptible, non-persister strains, but it may not be able to eliminate persister strains with the same efficiency in all individuals. We infected larvae with 10^4^ (Fig. 7A) or 5×10^4^ (Fig. 7B) conidia of non-persister (ATCC46645 or PD-60) or persister (PD-9 or PD-104) isolates, administered one dose of 8 µg/larva voriconazole or PBS at the time of infection, and measured fungal burden 72 hours (Fig. 7A) or 48 hours (Fig. 7B) after infection. As expected, voriconazole treatment was efficient and dramatically decreased fungal burden for all isolates (Fig. 7A and B). To note, for voriconazole treated groups, the mean of fungal burden was bigger for persisters than for non-persister isolates, although these differences were not significant (ATCC VS PD-9 *p*=0.41 for 10^4^ and *p*=0.57 for 5×10^4^, ATCC VS PD-104 *p*=0.27 for 10^4^ and *p*=0.69 for 5×10^4^, Mann-Whitney test), due to the high variability in burden among individuals infected with the persister isolates. Indeed, at both infectious doses several larvae infected with persister isolates had noticeable greater burdens than all the other (Fig. 7A and B), which we speculate might indicate that in some individuals infected with persistent isolates treatment may be less efficient in eliminating the fungus.

**Figure 7.**
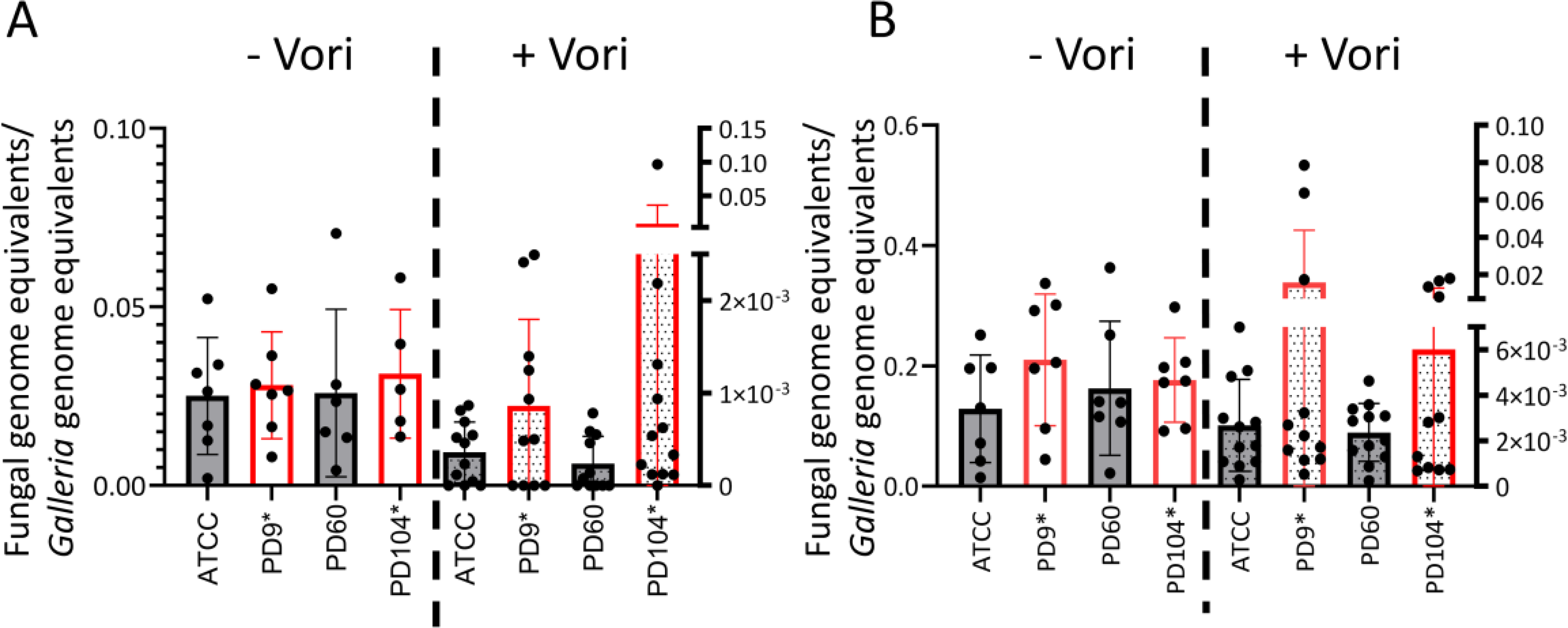
In some larvae, voriconazole eliminates persister isolates less efficiently than non-persister strains in a *Galleria mellonella* infection model. Fungal burden was measured by qPCR **A)** 72 h after infection with 10^4^ conidia, or at **B)** 48 h after infection with 5×10^4^ conidia For all strains, fungal burden was greatly lower in larvae that had received a voriconazole treatment (8 µg/mL) that in those who had not. The reduction in burden was bigger in non-persister isolates than in persister isolates, although these differences were not significant. Several treated larvae infected with persister isolates had noticeable higher burdens than the other: **A)** 4/11 for PD-9 and 3/11 for PD-104 **B)** 3/11 for PD-9 and 4/11 for PD-104. Each graph displays the combined data from two independent experiments.

## DISCUSSION

The phenomenon of persistence to antimicrobials was first observed in bacteria 80 years ago [78, 79]. In the last decade, intensive research has permitted unravelling various underlying mechanisms in diverse bacteria, and it has become clear that persistence can cause treatment failure and lead to the development of resistance [80, 81]. However, our insight into antifungal persistence and tolerance in fungal pathogens is still in its infancy.

We have recently shown that all conidia in a tolerant *A. fumigatus* isolate are able to grow at high concentrations of caspofungin [26]. In *C. albicans*, tolerance to fluconazole was described to be a sub- population effect, but the ratio of tolerant cells was reported to be elevated [21]. In both cases, tolerance was observed in response to static drugs, so it seems that the phenomenon of tolerance in fungi associates with fungistatic drugs. In *C. albicans*, the formation of amphotericin-B (AmB) persister cells has been described [82]. Of note, this phenomenon has only been described within biofilms, and cannot be found in planktonic cells [83], possibly due to the physical protection conferred by the biofilm, which may be relevant for the capacity of these cells to survive in the presence of AmB. Indeed, in *A. fumigatus* it was shown that the hypoxic microenvironment developed in biofilms led to reduced metabolic activity in the basal biofilm level leading to the formation of cells that can survive the antifungal challenge, and serve as a drug-resistant reservoir [84].

We have observed that a small sub-population (0.1 to 5%) of certain *A. fumigatus* isolates can survive for extended periods and even grow at slow rates in the presence of supra-MIC concentrations of the fungicidal drug voriconazole. In bacteria, tolerance and persistence have been classically explained by a downregulation of metabolism and cell cycle, which trigger a status of dormancy that permit survival despite the action of the drug. However, recent research in pathogenic bacteria has demonstrated that active metabolic responses are required for persistence [85, 86], and slow-growing persisters have also been detected *in vitro* [87] and *in vivo* [88]. Active metabolism may even be advantageous, as in the case of *Salmonella*, where persisters have been shown to undermine host defences [89]. Interestingly, *C. albicans* AmB persistence has been described to involve downregulation of primary metabolism, but also an increase of stress responses and oxidative defensive mechanisms [90]. We have observed that active growth is possible at two or even three-fold the MIC, and that this growth entails a distinct transcriptional profile. Therefore, this phenomenon is not merely survival of dormant conidia in the presence of the drug, but seems to be an active mechanism that enables a sub- population of certain isolates to withstand the action of voriconazole for an extended period. Indeed, using resazurin (an oxidation-reduction indicator used for the measurement of metabolic activity and proliferation of living cells, which use has been optimised for *A. fumigatus* [91]) we could detect a slight metabolic activity at supra-MIC concentrations (Fig. S9C). This might explain why hypoxia is the only environmental condition that reduced persistence, as in low oxygen the metabolic activity is reduced [84] and the energy generating metabolism changes [92].

We have observed a big difference in growth between liquid media (only microscopic growth) and solid media (macroscopic colonies). We hypothesise that this may be because in agar drug microdepletions around the hyphae would diffuse back much slower than in liquid, giving an advantage to the hyphae in solid media. In addition, there may be a different level of contact of conidia with the drug; in liquid, the conidia are completely surrounded, whilst on solid media the conidia just lay on the drug and may escape from it by orientating the germling polarity as they grow.

It is surprising that we have not found any voriconazole tolerant isolate. It might be that the frequency of this phenomenon is lower in *A. fumigatus* (even in filamentous fungi in general), and we simply have not screened enough isolates to detect it. However, we believe that the particularities of working with filamentous fungi make the definitions developed with bacteria unsuitable. Firstly, MIC measurements or killing curves need to be done over a period of days, compared with hours for bacteria. In addition, the inoculum used is normally conidia, a dormant cell structure that needs to be activated to produce an effect. Besides, the experiments are performed with inocula of around 10^4^ (2.5×10^4^ for broth dilution and 4×10^4^ for disc assays) conidia, whilst bacteria persistence can be assayed with inocula in the range of 10^6^ – 10^8^ cells. Therefore, we believe it would be challenging to clearly discern between the duration of killing for 99% or 99.99% of the cells, the most important factor to differentiate tolerance and persistence [18, 19]. Indeed, looking closely at the killing curve presented in Fig. 1D, at 2 days the percentage of survivors is 99.38% for PD-9 and 99.86% for PD-104. Could this mean that PD-9 is tolerant and PD-104 a persister? This seems to be supported by the experiments plating wells from the broth dilution assays (Fig. 1C and 5C), where more colonies are counted for PD-9 than for PD-104. However, the number of colonies detected in the disc assays is usually higher for PD-104 (2-5) than for PD-9 (1-2). Given that the mechanisms of tolerance and persistence in bacteria are multiple and overlapping [93], we propose to define the phenomenon that we have described in this study as persistence and utilize the term tolerance to define the effect previously described with fungistatic drugs [21, 26]. Therefore, we propose to adapt the current definitions for pathogenic fungi as presented in Table 1.

The concept of persistence entails two intriguing aspects. Firstly, it is an isolate-dependent phenomenon, which means that there must be a genetic basis that underlies persistence. We have shown that the persister isolates do not belong to a specific lineage, suggesting that this feature has not appeared as an evolutionary trait in a lineage of isolates. Recently, it has been described that each *A. fumigatus* strain carries a particular set of accessory genes, which associates with different levels of virulence and drug resistance capacities [94]. In the same vein, we speculate that the presence or absence of certain accessory genes may enable certain isolates to persist in the presence of azoles. Future research will evaluate if this hypothesis is true. A second intriguing aspect of persistence is that it is a sub-population phenomenon, meaning that within an isogenic isolate only a few conidia are able to survive and grow in the presence of the drug. In bacteria, stochastic expression of key genes has often been proposed as an important underlying cause of persistence [68, 69, 95, 96]. This includes stochastic high expression of genes that directly confer resistance [97] and efflux activity [98]. Similarly, in our RNA-seq analysis we have detected a higher level of expression of genes of the sterol biosynthesis pathway, including *cyp51A*, and of the azole exporter Cdr1B in persister cells. Therefore, it is plausible that stochastic high levels of these genes could generate the capacity to survive for an extended period in the presence of supra-MIC concentrations of voriconazole. Additionally, we detected a higher level of expression of GAG biosynthetic genes, and we have observed that externally added GAG can increase the number of persisters in the isolate PD-104. Interestingly, this polysaccharide did not increase the isolate’s MIC, indicating that it does not provide a physical barrier that shields the fungus or that it cannot somehow degrade the drug. Future investigations will aim to unravel why GAG potentiates persistence in PD-104, and not in PD-9, which is interesting as it suggests that there might be multiple persistence mechanisms. GAG is a very relevant *A. fumigatus* virulence factor, with multiple described adhesion and immunosuppressive activities [99, 100], and here we propose that it may have one more important role in potentiating persistence to voriconazole.

In bacterial infections, there is significant evidence supporting a role for the phenomena of tolerance and persistence in antibiotic treatment failure [16, 101]. In contrast, the knowledge about these phenomena in fungal infections is still very scarce. Probably, the best-studied phenomenon is heteroresistance in *Cryptococcus neoformans* and *C. gatii* [102–104], which has been found to be a major cause of treatment failure in cryptococcal meningitis [105–109]. As mentioned above, there is already evidence indicating that fluconazole tolerance can cause treatment failure in invasive candidiasis [17, 21], and limited evidence suggests that caspofungin tolerance may cause treatment failure in *Aspergillus* infections [27]. Here, using a *Galleria* model of infection, we have observed that voriconazole treatment seems to eliminate persister isolates less efficiently than non-persister isolates in various larvae. Therefore, we hypothesise that in certain individuals persistence might reduce the efficacy of voriconazole treatment in *A. fumigatus* infections, which, if proven true, could imply that persistence may cause therapeutic failure in some patients. In addition to such a potential direct effect of persistence on treatment failure, resilience and relapse of persistent strains could associate with the development of antifungal resistance (an effect that has already been shown for bacterial infections, see [110] for references), as it is known that prolonged azole treatment can result in resistance [111, 112]. We appreciate that our results provide limited evidence to support the hypothesis that voriconazole persistence may cause treatment failure, and acknowledge that much more research is required to reach such conclusion. We will continue investigating this hypothesis and we exhort the fungal research community to consider and investigate this phenomenon to clarify this important matter.

## MATERIAL and METHODS

### *A. fumigatus* strains and culture conditions

All isolates utilized in the course of this study are listed in Tables S1 and S8. Briefly, information about the common laboratory strains used can be found in [113], the isolates from Paul Dyer collection are described in [114] and the collection of isolates from the area of Manchester are described in [73]. The third collection of isolates was collected at the TAU medical centre (Tel-Aviv, Israel), these strains were characterized for their antifungal resistance profile and *cyp51A* genotype. Persister isolates are labelled in all figures with an asterisk (*).

Isolates were routinely grown on Potato Dextrose Agar (PDA, Oxoid) for 72 hours to obtain fresh spores for each experiment. All experiments were performed with fresh spores except in the test for persistence with old conidia. *Aspergillus* Minimal Medium (AMM) was prepared following a standard recipe [115]. Sabouraud (Oxoid) and yeast extract glucose (YAG, 2%, 0.5% yeast extract, 1.7% agar, 1X trace elements) media were used in specific experiments.

To evaluate persister growth and determine the MIC, isolates were grown on RPMI-1640 (Sigma) with 35 g/L MOPS (Alfa Aesar) and 2% glucose, pH 7.

Galactosaminogalactan (GAG) was obtained from the *A. fumigatus* Δ*ku80* strain, grown 2-day in 1.5L brian fermenter at room temperature. GAG isolation and purification was carried out as previously described [116]. Briefly, the medium supernatant was collected by filtration and was adjusted to pH 3 by addition of 100 μl 12 M HCl per 100 ml supernatant. Two volumes of precooled ethanol (4°C) were added and GAG was precipitated for 3 h at 4°C. The precipitate was collected by centrifugation for 20 min at 5,000 g at 4°C and subsequently washed twice with 1/10 of the culture volume of 200 mM NaCl for 1 h under agitation (100 rpm). GAG was dialyzed against tap water and twice against purified water (24 h each) and finally lyophilized to dryness and stored at ambient temperature.

To calculate germination rate, 10^4^ conidia of each strain were inoculated in 200 µL liquid RPMI in 96- well plates and imaged a Nikon TI microscope equipped with a 37 °C incubator and a 40× objective, with 1 picture captured every 30 min by using NIS-Elements 4.0 (Nikon) software. Cell Counter plug- in of Image J platform (http://rsb.info.nih.gov/ij/index.html) was employed to differentially count resting or swollen conidia versus germinated conidia.

To calculate growth rate on solid medium, 10^3^ conidia of each strain were inoculated on RPMI plates, and the colony diameter was measured in two different angles per colony at 16, 24, 40 and 48 h. To calculate growth rate on liquid medium 2×10^3^ of each strain were inoculated in 200 µL liquid RPMI in 96-well plates and optical density measurements were taken at 600 nm every 10 minutes. Growth curves were analysed using the R package Bayesian Estimation of Change-points in the Slope of Multivariate Time-Series (BEAST) [117] With parameters; burn = 2000, 5000 iterations, zeroNormalization=TRUE and a blankThreshold of 0.095. Confidence intervals of the first change-point were plotted in Graphpad PRISM.

### Evaluation of persistence and determination of MICs

To determine persistence using the disc assay, 4×10^4^ conidia of each isolate was evenly spread on a solidified RPMI plate (1.5% agar) and 10 μl of 0.8 mg/mL voriconazole or 3.2 mg/mL itraconazole (Acros Organics) was added to a Whatman 6 mm antibiotic assay disc which was placed in the middle of the plate. For *in vitro* treatment with adjuvant and combinatorial drugs, the agents were added to the RPMI medium at the final concentration detailed in each section. Plates were incubated for 5 days at 37 °C. Persister colonies were defined as those with a degree of physical separation from the edge of the loan of growth into the halo of inhibition.

Minimum inhibitory concentrations (MICs) were calculated using the broth microdilution method according to the EUCAST E.Def. 9.3 instructions [118]. 2.5×10^4^ conidia were used in each well.

### Evaluation of the conidial survival to voriconazole fungicidal action

To calculate the number of conidia that survived after exposure to high concentration of azoles, full contents of the wells containing the highest concentration of each drug (8 µg/mL) in the broth dilution assays were plated after 48-72 hours incubation on PDA and further incubated for 24-48 h at 37 °C.

Microscopic analysis was used to investigate the survival of strains upon withdrawal of voriconazole at single cell level. 1.5×10^4^ conidia of each strain were inoculated in 300 µL of RPMI in the presence of 8 µg/mL voriconazole (European Pharmacopoeia, -EP- Reference Standard, Merck) and incubated for 48 hours at 37 °C with occasional shaking. The strains were then centrifuged for 4000 RPM for 5 minutes, the media discarded and the conidia resuspended in 2 mL filtered PBS. This wash was repeated three times in total before the conidia were resuspended in 600 µL of drug free RPMI. 300 µL of each suspension, approximately 7.5 ×10^3^ conidia, was added into wells of a 96 well plate. The wells were then imaged for 72 hours, every 2 hours, on a Nikon Eclipse Ti microscope, using a Nikon CFI Plan Fluor ELWD 20x/0.45na objective, and captured with a Hamamatsu ORCA-FLASH4.0 LT+ camera (Hamamatsu Photonics) and manipulated using NIS-Elements AR 5.11.01 (Nikon). A video was prepared using Fiji [119].

Microscopy to observe growth at supra-MIC conditions was done in a THUNDER Imager Live Cell microscope, with a HC PL FLUOTAR L 20x/0.40 DRY objective. Images were captured using a Leica- DFC9000GTC-VSC13067 camera and the Las X (Leica application suite) v 3.7 software.

To construct the killing curved, the isolates were inoculated in 10 mL of RPMI containing 4 µg/mL voriconazole. Aliquots for each culture were taken at the time of inoculation (100 µL) and every 24 hours (1 mL). The aliquots were spun at 16,000 ×g for 5 minutes, resupended in 1 mL PBS and vortexed. This was repeated twice to wash off the drug. Finally, conidia were resuspended in 1 mL PBS and a fraction plated on PDA plates (50 µL at time 0h, 100 µL at 24h and 1 mL for 48, 72 and 96 h). The experiment was repeated three times.

To calculate the percentage of germinating conidia, the wells of broth dilution assays containing 2X, 3X MIC and the maximum drug concentration (8 µg/mL) were stained (after reading the MIC) with 10 µg/mL of Calcofluor White (Sigma) for 5 minutes. The entire wells were imaged with a THUNDER

Imager Live Cell microscope, using a HC PL FLUOTAR L 20x/0.40 DRY objective, filter conditions: EX:375-435 DCC:455 EM:450-490, a Leica-DFC9000GTC-VSC13067 camera and the Las X (Leica application suite) v 3.7 software. Merged images were analysed using FIJI [119]. Briefly, the merged images were converted to 8-bit, a threshold was set for the images so that the conidia and germlings could be detected over the background (20 to 255). The option analyse particle was then executed, setting a minimum size of 0.04 inches^2^ (which was found to exclude resting conidia). Wells containing the maximum concentration of the drug, where only resting conidia can be found, were used to set the background level of detection for each isolate (consisting of aggregates of conidia, impurities and carry over conidiophores from the inoculum). The percentage of germinated conidia/short hyphae was calculated as: (number of counted particles per well – number of counted particles in max drug)/25000 (inoculum).

### Nucleic acid isolation

For DNA extraction, 10^6^ spores of each *A. fumigatus* isolate were grown overnight on liquid *Aspergillus* minimal medium, filtered through Miracloth paper (Merck), snap frozen and ground in constant presence of liquid nitrogen. Subsequently, DNA was extracted using a standard CTAB method.

For RNA extraction, *A. fumigatus* isolates were grown on 0.45 µm pore nylon membranes (GE Healthcare) placed on RPMI plates and incubated 1 day (without drug) or 5 days (with voriconazole in a disc). Mycelia were scratched using a spatula and immediately frozen and grounded in liquid nitrogen. Subsequently, RNA was extracted and purified with the Plant RNeasy Mini Kit (Qiagen) following the manufacturer’s instructions for filamentous fungi, and using on-column DNase treatment.

For fungal burden calculations in *Galleria mellonella*, DNA was extracted from decapitated larvae using the DNeasy Blood & Tissue Kit (Qiagen) following the manufacturer’s instruction for animal tissue, with overnight incubation at 56 °C.

### *Galleria mellonella* infection

Sixth-stage instar larval *G. mellonella* moths were purchased from R.J.Mous Livebait (Eigen Haard, The Netherlands). Randomly selected groups of 250-350 mg of weight larvae were injected in the last left proleg with 5 µL of a 2×10^6^ conidia/mL suspension (10^4^ conidia/larva) of the correspondent *Aspergillus fumigatus* isolate using Hamilton syringe. 2 hours after infection, relevant groups were injected in the last right proleg with 5 µL of a suspension containing 1600 µg/mL (8 µg/larvae) voriconazole. A PBS control group was subjected to the same treatment, but without fungal infection.

To detect the fungal burden, 500-ng portions of DNA extracted from each larva were subjected to qPCR using the SensiMic SybR Green kit (Bioline). Forward (5′-ACTTCCGCAATGGACGTTAC-3′) and reverse (5′-GGATGTTGTTGGGAATCCAC-3’) primers were used to amplify the *A. fumigatus* β-tubulin gene (AFUA_7G00250). The primers designed to amplify the Elongation factor 1-Alpha (Ef-1a) were as follows: forward (5′- AACCTCCTTACAGTGAATCC-3′) and reverse (5′- ATGTTATCTCCGTGCCAG-3′). Standard curves were calculated using different concentrations of fungal and larval gDNA pure template. Negative controls containing no template DNA were subjected to the same procedure to exclude or detect any possible contamination. Three technical replicates were prepared for each sample. qPCRs were performed using a CFX96 Real-Time System (Bio-rad) with the following thermal cycling parameters: 95°C for 10 min and 40 cycles of 95°C for 15 s and 58°C for 15 s and 72°C for 15 s. The fungal burden was calculated by normalizing the number of fungal genome equivalents (i.e., the number of copies of the tubulin gene) to the larval genome equivalents in the sample (i.e., the number of copies of the a Ef-1a gene), as we have reported before [120].

### RNA sequencing (RNA-seq)

Total RNA was submitted to the Genomic Technologies Core Facility (GTCF) at the University of Manchester, UK. Quality and integrity of the RNA samples were assessed using a 4200 TapeStation (Agilent Technologies) and then libraries generated using the Illumina® Stranded mRNA Prep. Ligation kit (Illumina, Inc.) according to the manufacturer’s protocol. Briefly, total RNA (typically 0.025-1ug) was used as input material from which polyadenylated mRNA was purified using poly-T, oligo-attached, magnetic beads. Next, the mRNA was fragmented under elevated temperature and then reverse transcribed into first strand cDNA using random hexamer primers and in the presence of Actinomycin D (thus improving strand specificity whilst mitigating spurious DNA-dependent synthesis). Following removal of the template RNA, second strand cDNA was then synthesized to yield blunt-ended, double-stranded cDNA fragments. Strand specificity was maintained by the incorporation of deoxyuridine triphosphate (dUTP) in place of dTTP to quench the second strand during subsequent amplification. Following a single adenine (A) base addition, adapters with a corresponding, complementary thymine (T) overhang were ligated to the cDNA fragments. Pre-index anchors were then ligated to the ends of the double-stranded cDNA fragments to prepare them for dual indexing. A subsequent PCR amplification step was then used to add the index adapter sequences to create the final cDNA library. The adapter indices enabled the multiplexing of the libraries, which were pooled prior to cluster generation using a cBot instrument. The loaded flow-cell was then paired-end sequenced (76 + 76 cycles, plus indices) on an Illumina HiSeq4000 instrument. Finally, the output data was demultiplexed and BCL-to-Fastq conversion performed using Illumina’s bcl2fastq software, version 2.20.0.422

### RNA-seq analysis

Unmapped paired-reads of 76bp from the Illumina HiSeq4000 sequencer were checked using a quality control pipeline consisting of FastQC v0.11.3 (http://www.bioinformatics.babraham.ac.uk/projects/fastqc/) and FastQ Screen v0.13.0 (https://www.bioinformatics.babraham.ac.uk/projects/fastq_screen/).

The reads were trimmed to remove any adapter or poor quality sequence using Trimmomatic v0.39 [121]; reads were truncated at a sliding 4bp window, starting 5’, with a mean quality <Q20, and removed if the final length was less than 35bp. Additional flags included: ’ILLUMINACLIP:./Truseq3- PE-2_Nextera-PE.fa:2:30:10 SLIDINGWINDOW:4:20 MINLEN:35’.

The filtered reads were mapped either to the *Aspergillus fumigatus* A1163 reference sequence (GCA_000150145.1/ASM15014v1) downloaded from the Ensembl Genomes Fungi v44 [122] or PD- 104 a *de novo* assembled genome, using STAR v5.3a [123]. For A1163 reference the genome index was created using the GTF gene annotation also from Ensembl Genomes Fungi v44. For the PD-104 reference GTF gene annotation was generated as described below. A suitable flag for the read length (--sjdbOverhang 75) was used. ’--quantMode GeneCounts’ was used to generate read counts in genes.

Subsequently, PD-104 reads were aligned to the recently generated pan-genome [65] using Salmon v1.6.0 (additional parameters for salmon quant: --gcBias (corrects for any GC biases in samples) -l ISR (inward / stranded / reverse read pairs)).

Normalisation and differential expression analysis was performed using DESeq2 v1.34.0 [124] on R v4.1.2 (R Core Team. 2021. R: A Language and Environment for Statistical Computing. https://www.R-project.org/). Log fold change shrinkage was applied using the lfcShrink function along with the “apeglm” algorithm [125].

The RNA data is stored in Array Express, accession E-MTAB-11547 (https://www.ebi.ac.uk/arrayexpress/experiments/E-MTAB-11547).

### DNA sequencing (Illumina DNA Prep)

Genomic DNA was submitted to the Genomic Technologies Core Facility (GTCF) at the University of Manchester. Sequencing libraries were generated using on-bead tagmentation chemistry with the llumina® DNA Prep, (M) Tagmentation Kit (Illumina, Inc.) according to the manufacturer’s protocol. Briefly, bead-linked transposomes were used to mediate the simultaneous fragmentation of gDNA (100-500ng) and the addition of Illumina sequencing primers. Next, reduced-cycle PCR amplification was used to amplify sequencing ready DNA fragments and to add the indices and adapters. Finally, sequencing-ready fragments were washed and pooled prior to paired-end sequencing (151 + 151 cycles, plus indices) on an Illumina NextSeq500 instrument. Finally, the output data were demultiplexed (allowing one mismatch) and BCL-to-Fastq conversion performed using Illumina’s bcl2fastq software, version 2.20.0.422.

Method for de novo genome assembly of genomes (PD-104 as example):

Unmapped paired-end reads of 149bp from an Illumina NextSeq500 sequencer were checked using a quality control pipeline consisting of FastQC v0.11.3 (http://www.bioinformatics.babraham.ac.uk/projects/fastqc/) and FastQ Screen v0.13.0 (https://www.bioinformatics.babraham.ac.uk/projects/fastq_screen/).

The reads were trimmed to remove any adapter or poor quality sequence using Trimmomatic v0.39 [121]. Within the first or last 30bp of a read bases were removed if quality was <30. A sliding 2bp window, starting 5’, scanned the reads and truncated at a window with a mean quality <Q20. Reads were removed if the final length was less than 100bp. Additional flags included: ’ILLUMINACLIP:./Truseq3_Nextera-PE.fa:2:30:10 LEADING:30 TRAILING:30 SLIDINGWINDOW:2:20 MINLEN:100’.

Genome assembly was performed using Megahit v1.2.9 [126], using the flag ’--no-mercy’. This resulted in 799 contigs, total 28659847 bp, min 202 bp, max 401152 bp, avg 35869 bp, N50 97740 bp. In comparison A1163 has 55 contigs total 29205420 bp.

Quast v5.0.2 [127] was used to determine the quality of the assembly. An example of the PD-104 results can be found in Table S10 in https://doi.org/10.5281/zenodo.7021623.

ProtHint v2.5.0 and GeneMark-EP+ v4.68_lic [128] were used for gene annotation. ProtHint was used to generate the evidence for coding genes in combination with fungal protein download from OrthoDb v10. ProtHint generates two main outputs: ‘prothint.gff’ a GFF file with all reported hints (introns, starts and stops), and ‘evidence.gff’ high confidence subset of ‘prothint.gff’ which is suitable for the GeneMark-EP Plus mode. GeneMark-EP+ was used to generate a GTF file of candidate genes for the PD-104 assembly. Additional parameters were set ’--max_intron 200 --max_intergenic 100000 -- min_contig=1000 --fungus’.

In order to determine the function of the novel genes, initially a reciprocal best hit strategy was used comparing protein sequence from PD-104 (using the GeneMark Perl script ’get_sequence_from_GTF.pl’ with the GTF file and assembled contigs) and A1163 (proteins downloaded from Ensembl Genomes Fungi release 51: http://ftp.ensemblgenomes.org/pub/fungi/release-51/fasta/aspergillus_fumigatusa1163/pep/Aspergillus_fumigatusa1163.ASM15014v1.pep.all.fa.gz). The best match method was run using proteinortho v6.0.31 [129], which by default uses the program Diamond v2.0.13 [130] to identify protein matches (’-p=diamond’). The flag ’-sim=1’ was set to obtain best reciprocal matches only.

### Construction of Phylogenetic tree

Pre-alignment, the reference Af293 genome [131] was masked to remove low-complexity regions using RepeatMasker (http://www.repeatmasker.org/) v.4.1.2-p1 and the Dfam repeat database [132] release 3.5. Quality-checked DNA sequencing reads from the recently generated pan-genome study [65] were combined with the reads generated from this study and underwent alignment to the masked reference Af293 genome, using BWA-MEM [133] v.0.7.17-r1188. Alignment files were then converted to the sorted BAM format using SAMtools [134] v.1.10. Variant calling was then performed with GATK HaplotypeCaller [135] v.4.0. All variant calls with a genotype quality less than 50 were removed and low-confidence SNPs were converted to missing data if they met one of the follow criteria: DP < 10 || RMSMappingQuality < 40.0 || QualByDepth < 2.0 || FisherStrand > 60.0 || ABHom < 0.9. Each newly generated VCF file was then compressed, indexed and then merged into a single file using VCFtools [136] v.0.1.16. This data was then converted into a relaxed interleaved Phylip format using vcf2phylip [137] v2.8 and RAxML [138] v.8.2.12, under rapid bootstrap analysis using 1,000 replicates with the GTRCAT model, was employed to generate maximum-likelihood phylogenies to test whether persistence is lineage-specific. Phylogenetic trees with overlaying metadata were generated using iTOL [139] v.6.5.4.

## Supporting information

DatSet S1

DatSet S2

DatSet S4

## ACKNOWLEGDGEMENTS

We are deeply grateful to Prof Paul Dyer, who kindly gifted to us the collection of isolates that have been used in this study. We would like to thank Diego Megías and Clara Martín at ISCIII for their help with the microscopy. We are also grateful to Michael Bottery for relevant conversations about bacterial tolerance and persistence. We acknowledge the use of the Genomic Technologies Facility (Faculty of Biology Medicine and Health, University of Manchester) for the DNA and RNA sequencing. Help and support from members of MFIG and LRIM is greatly appreciated.

JA is funded by an Atracción de Talento Modalidad 1 (020-T1/BMD-200) contract of the Madrid Regional Government (CAM). JS has been funded by a BSAC Scholarship (bsac-2016-0049). CV was funded by FAPESP (2108/00715-3 and 2020/01131-5). GHG has been funded by FAPESP (2016/07870- 9 and 2021/04977-5), CNPq (301058/2019-9 and 404735/2018-5) and by the NIH/NIAID (grant no. R01AI153356). SG was co-funded by the NIHR Manchester Research Centre and the Fungal Infection Trust.

## DATA AVAILABILITY

Transcriptomic data is available in Array Express (https://www.ebi.ac.uk/arrayexpress/experiments/E-MTAB-11547). All raw data is available upon reasonable request.

## Supplementary Figures

**Fig. S1.**

Disc diffusion screening of the PD-47 collection of isolates (Table S1 in https://doi.org/10.5281/zenodo.7021623) on RPMI plate with 10 µL of 0.8 mg/mL voriconazole. Only 5 isolates could grow colonies in the inhibition halos. Upon re-inoculation of the colonies of the halo (grown in the absence or presence of voriconazole, as detailed), PD-254, PD-256 and PD-266 covered the entire plate. As explained in the text those strains were classified as resistant or heteroresistant. In contrast, PD-9 and PD-104 CoHs showed upon re-inoculation the same level of growth as the original isolates. These strains were classified as persisters.

**Fig. S2.**

**A)** The standardized E-test strip provided the same results as disc diffusion assays: the persister isolates PD-9 and PD-104 developed colonies in the halo of inhibition, and the non-persister isolates ATCC46645 and PD-60 did not. E-test was performed according to Biomerieux instructions and repeated three independent times. **B)** Two CoHs from PD-9 and PD-104 were picked and sequentially passaged on PDA without drug. Every second passage the spores were inoculated in a new disc diffusion assay to check the level of resistance (diameter of the halo) and persistence (appearance of colonies in the halo). After 7 passages both CoHs from both isolates showed the same level of persistence, demonstrating that this phenotype is stable.

**Fig. S3.**

Disc diffusion screening of the PD-47 collection of isolates. Three days after inoculation, the disc containing voriconazole (10 µL of 0.8 mg/mL) was substituted for another one containing *Aspergillus* minimal medium. A few colonies were able to grow colonies in the halos after the switch, suggesting that a few conidia remain viable. Only three strains PD-9 and PD-104 and PD-259 grew colonies before the switch. Of those, only PD-9 and PD-104 grew additional colonies after the switch.

**Fig. S4.**

Independent repetitions of the experiment shown in Figure 1B. The wells of 1X, 2X and 3X MIC of voriconazole for the isolates ATCC, PD-9, PD-60 and PD-104 were imaged under the microscope. The non-persister isolates ATCC and PD-60 only displayed limited microscopic growth at 1X MIC. In contrast, the persister isolates PD-9 and PD-104 displayed microscopic growth at 1X, 2X and 3X supra-MIC concentrations. Scale bar= 132.5 µm.

**Fig. S5**

A) Photos taken at the site of ATCC conidia inoculation on an RPMI agar plate showed that germlings are visible after 8h of incubation at 37 °C, and short hyphae have developed after 16 hours of incubation. Agar was cut out an inverted on a cover slip containing a drop of PBS. Images were taken on a Leica SP8 inverted microscope. Images were processed using Fiji [119]. **B)** There was no difference in the germination rate or ratio of the isolates ATCC, PD-9, PD-60 and PD-104. **C)** Measurement of growth rate of persister and non-persister isolates on solid RPMI showed that the strains have no difference in basal growth. **D)** Measurement of growth rate of persister and non-persister isolates on liquid RPMI showed that the strains have no difference in basal growth.

**Fig. S6.**

Disc diffusion screening of the PD-47 collection of isolates (Table S1 https://doi.org/10.5281/zenodo.7021623) on RPMI plate with 10 µL of 3.2 mg/mL itraconazole. Isolates RC (as expected) and PD-256 did not form an inhibition halo, reflecting that they are resistant to itraconazole. Isolates PD-104 and PD-266 could grow colonies in the halo. PD-104 formed upon re- inoculation a halo of the same size and similar number of CoHs as the original isolate, suggesting that it is persistence to itraconazole. The isolate PD-266 nearly covered the inhibition halo upon reinoculation, suggesting an increment in its MIC and therefore that it is likely heteroresistant to itraconazole, as previously described for voriconazole.

**Fig. S7.**

**A)** MICs of itraconazole and isavuconazole for the isolates ATCC, PD-9, PD-60 and PD-104, as calculated by broth dilution assay. **B)** The isolates PD-9 and PD-104 displayed some microscopic growth at 2X the MIC concentration for both azoles, whilst the isolates ATCC and PD-60 did not grow at all and only resting conidia could be found. **C)** The entire content of the wells containing the highest concentration (8 µg/mL) if each azole were inoculated on PDA plates and incubated for 48 h at 37 °C. A noticeable ratio of viable conidia were detected for the isolates PD-9 and PD-104. Each experiment was repeated twice independently. The graph represents the means and SD

**Fig. S8.**

**A)** A nylon membrane was placed on the RPMI plate. Conidia were inoculated on the membrane and the disc containing voriconazole (10 µL of 0.8 mg/mL) was also put on it. As in a normal disc diffusion assay, an inhibition halo was formed, inside of which the persister strain PD-104 was able to grow small colonies, but the non-persister strain A1160 was not. **B)** A network of interacting proteins was detected by STRING (*p*=1.84e-08) when using as query the 64 genes upregulated only in persistence in PD-104. Seventeen proteins related with metabolism formed a node, formed a node suggesting a specific metabolic response during persister growth. **C)** Within the 18 genes that were upregulated in all comparisons, a tight node of 6 interacting proteins, related with ergosterol production, was detected by STRING (*p*<1.0e-16).

**Fig. S9.**

**A)** Representative disc diffusion plates carried out with the collection of isolates from the TAU medical centre. Six isolates were able to form CoHs, which did upon reinoculation form halos of the same size (MIC did not increase) and create a similar number of CoHs. **B)** Survival curves of larvae infected with 10^4^ or 5×10^4^ conidia of the isolates ATCC, PD-60 (non-persisters), PD-9 or PD-104 (persisters). All strains killed larvae at a similar rate, demonstrating that they are equally virulent. For 10^4^, three independent experiments were done with 15-20 larvae/isolate in each. For 5×10^4^, two independent experiments were done with 10-15 larvae/isolate in each. **C)** Resazurin was added to the broth dilution RPMI plate at a 0.002% (w/v) final concentration. The plate was incubated for 24 hours in a Tecan Infinity M-Plex plate reader and fluorescence was measured every 30 minutes with an excitation wavelength of 544 nm and reading emission at 590 nm using the i-control 2.0 software. The read for each well at each time point was normalize as follows: (read well time X– read 8 µg/mL voriconazole time X)/read well at time 0. The time point with the best dynamic range of values in the 1X, 2X and 3X MIC was found to be 20 hours, which was selected to determine the metabolic activity. A value of 0.1 was assigned as background and values above 1 were detected in sub-MIC (macroscopic growth). At 1X MIC, a slight metabolic activity could be detected for both the non-persister ATCC and the persister isolates PD-9 and PD-104. At 2X and 3X the MIC, slight metabolic activity could only be detected for the persister isolates. The experiment was performed once with two biological replicates, the graph represent the means and SEM.

## Supplementary Files (in https://doi.org/10.5281/zenodo.7021623)

**Table S1:** Collection of isolates used in this study.

**Table S2:** Synonymous polymorphisms detected in the *cyp51B* gene of the isolates ATCC46645, PD-9, PD-60 and PD-104.

**Table S3:** GO enrichment of genes regulated exclusively in Persistence (A1163 genome as reference).

**Table S4:** GO enrichment of genes regulated in Persiter VS Low Drug (Pangenome as reference).

**Table S5:** GO enrichment of genes regulated exclusively in Persistence (Pangenome as reference).

**Table S6:** Proteins included in the STRING network (genes upregulated only in Persistence).

**Table S7:** GO enrichment of genes regulated more in Persistence (Pangenome as reference).

**Table S8:** Proteins included in the STRING network (genes upregulated more in Persistence).

**Table S9:** List of environmental isolates in the UK collection. Identity of the sequenced genomes. List of the clinical isolates of the TAU collection.

**Table S10:** Example of the Quast obtained to determine the quality of the assembly, using the PD- 104 genome.

**DataSet1:** RNA-seq results and analysis of the A1160 isolate grown in NoDrug and Low Drug.

**DataSet2:** RNA-seq results and analysis of the PD-104 isolate grown in NoDrug, Low Drug and Persister, using the A1163 genome as reference for the alignment of reads.

**DataSet3:** RNA-seq results and analysis of the PD-104 isolate grown in NoDrug, Low Drug and Persister, using the pangenome genome as reference for the alignment of reads.

**Video S1:** The persister isolate PD-9 pre-incubated in voriconazole resumes growth when the drug is withdrawn.

**Video S2** The persister isolate PD-104 pre-incubated in voriconazole resumes growth when the drug is withdrawn.

**Video S3** The non-persister isolate ATCC46645 pre-incubated in voriconazole resumes growth when the drug is withdrawn.

