## Supplementary figures and images for "*Aspergillus fumigatus* can display persistence to the fungicidal drug voriconazole"

### Supplementary Figures

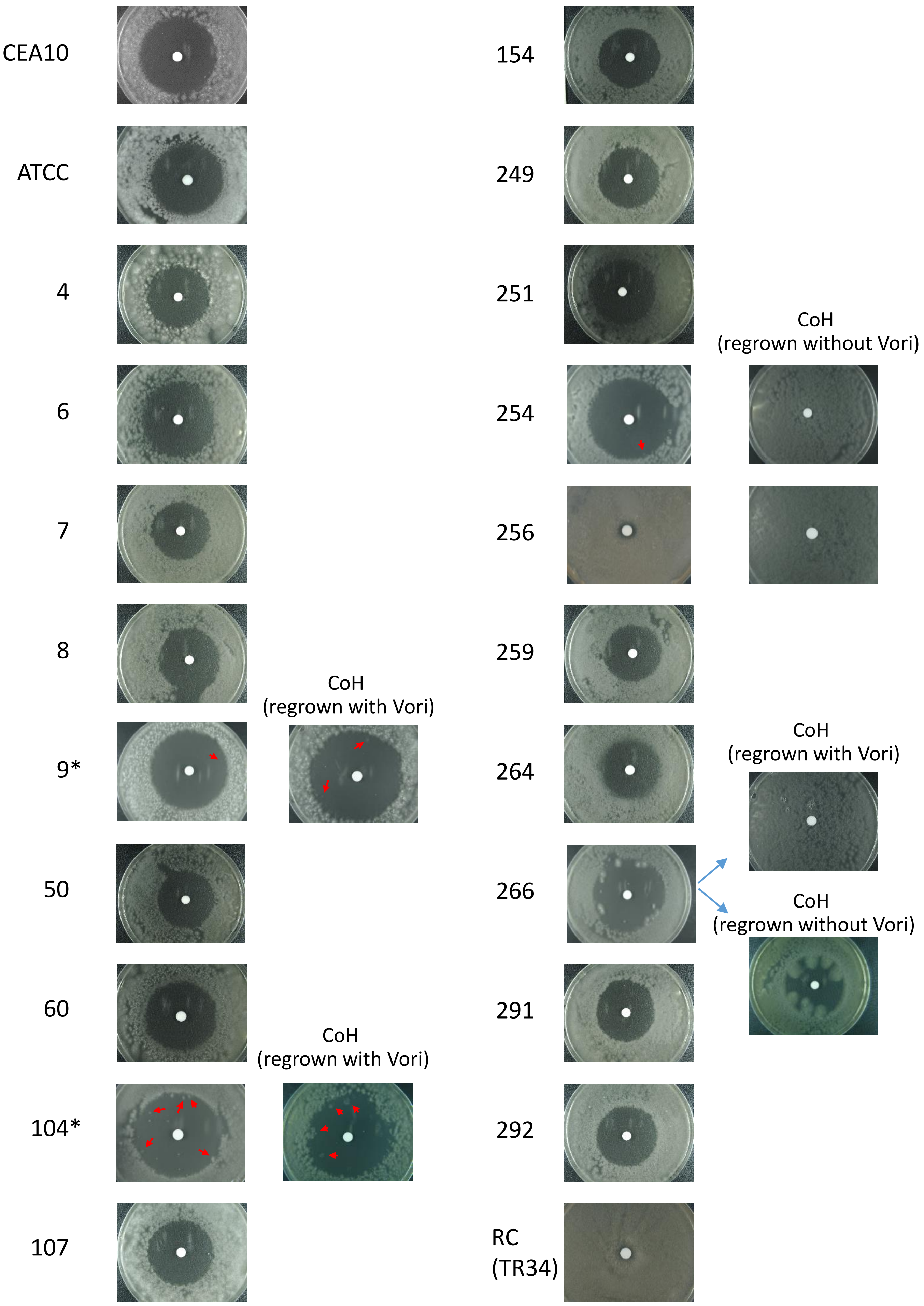

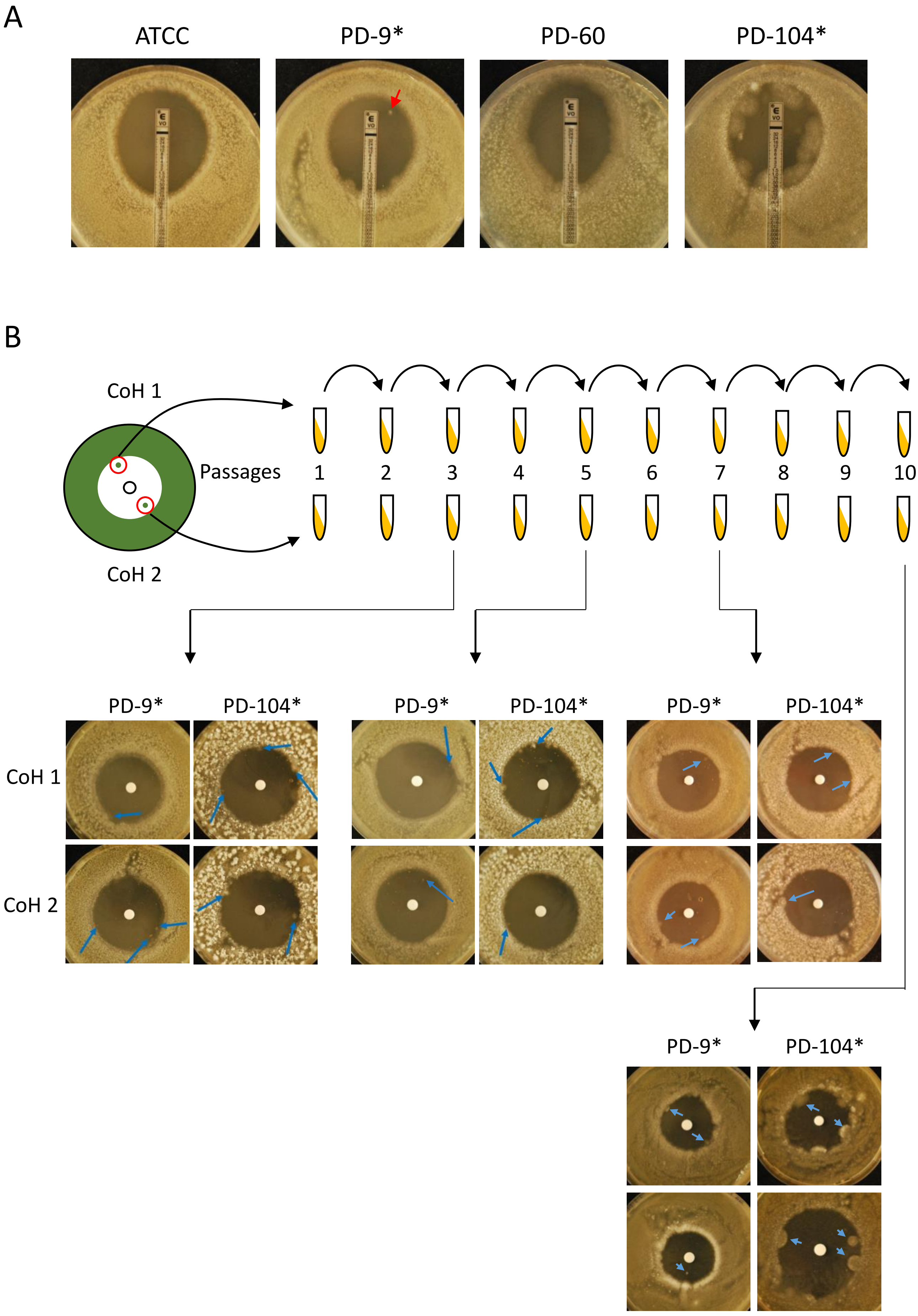

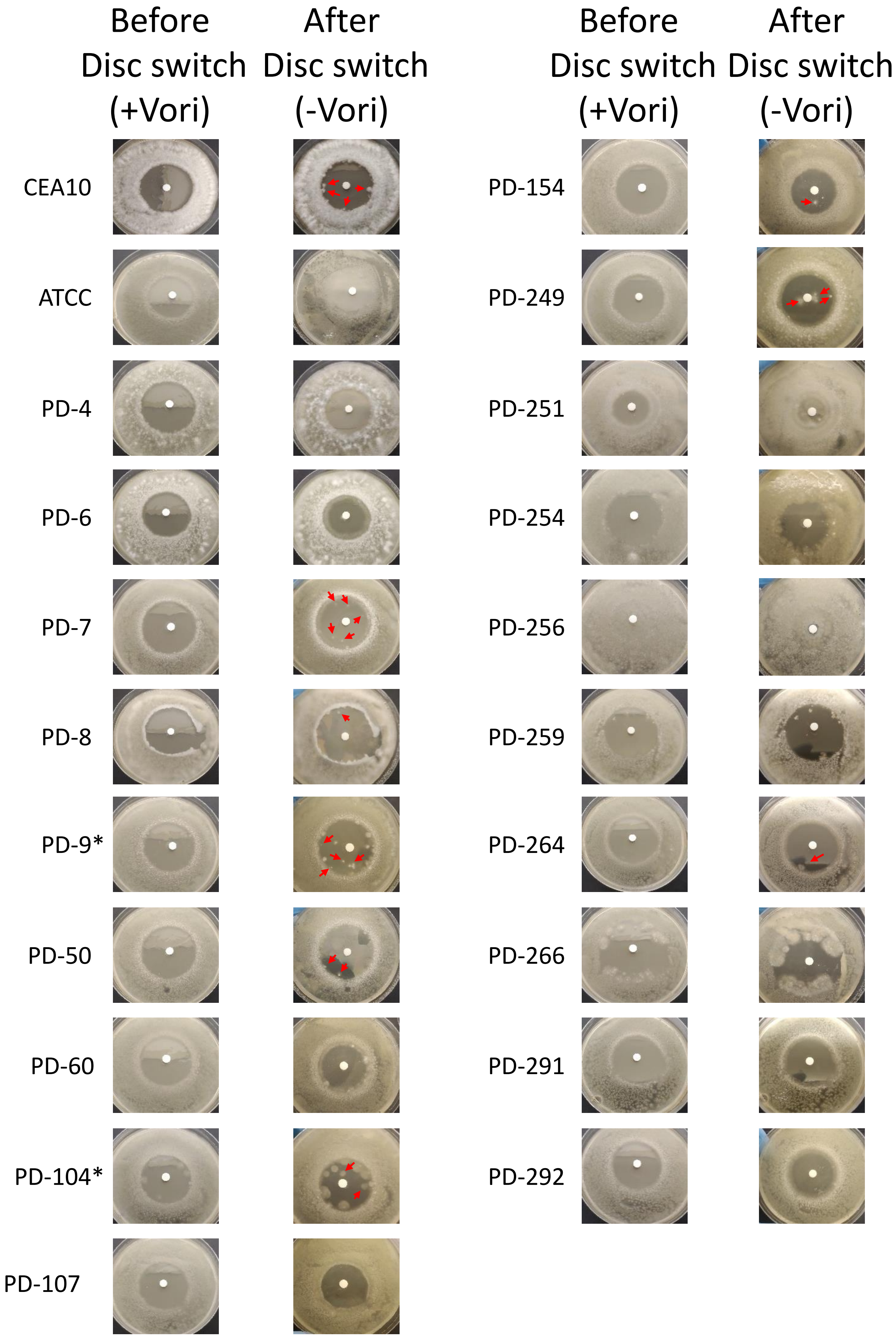

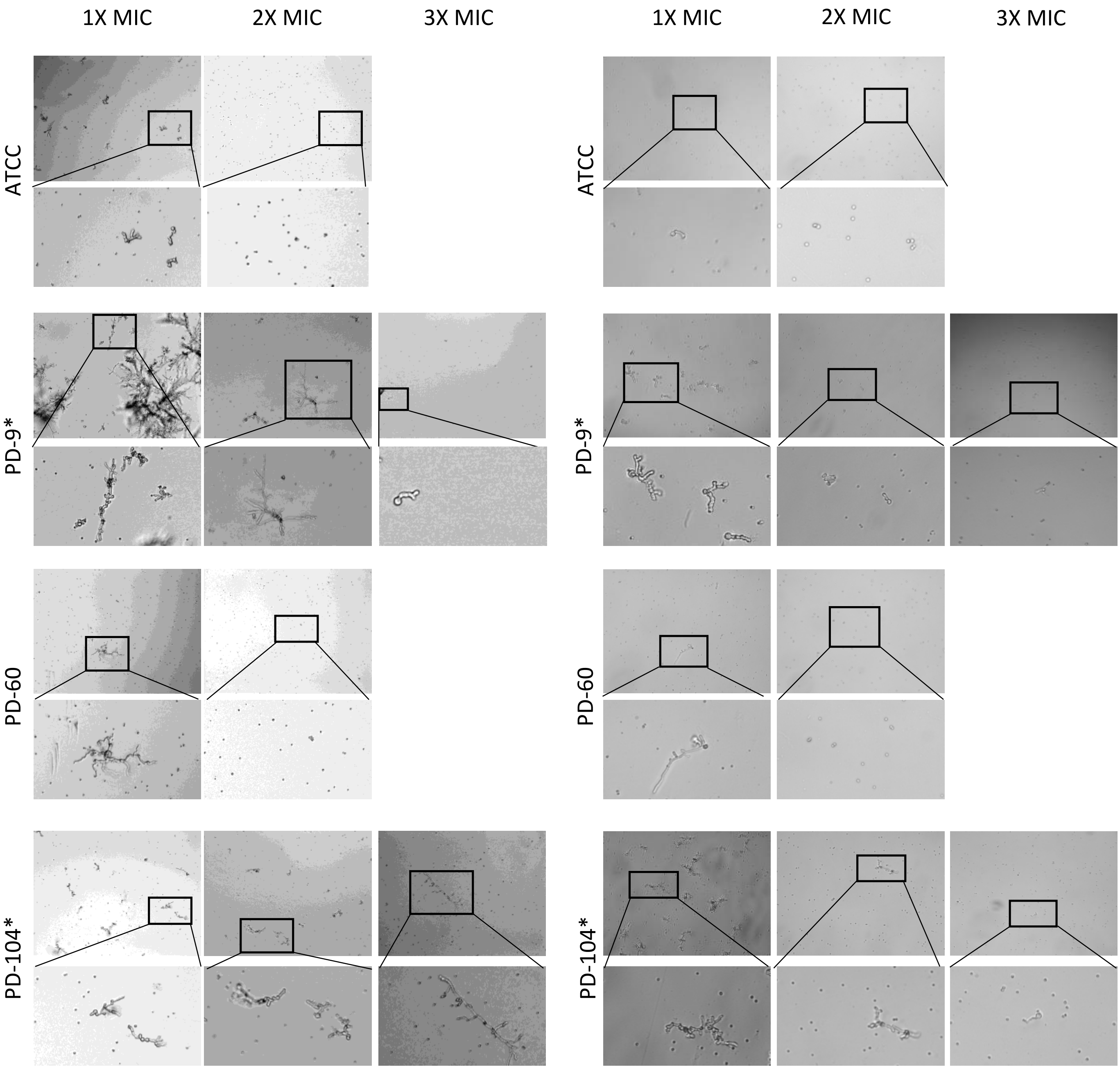

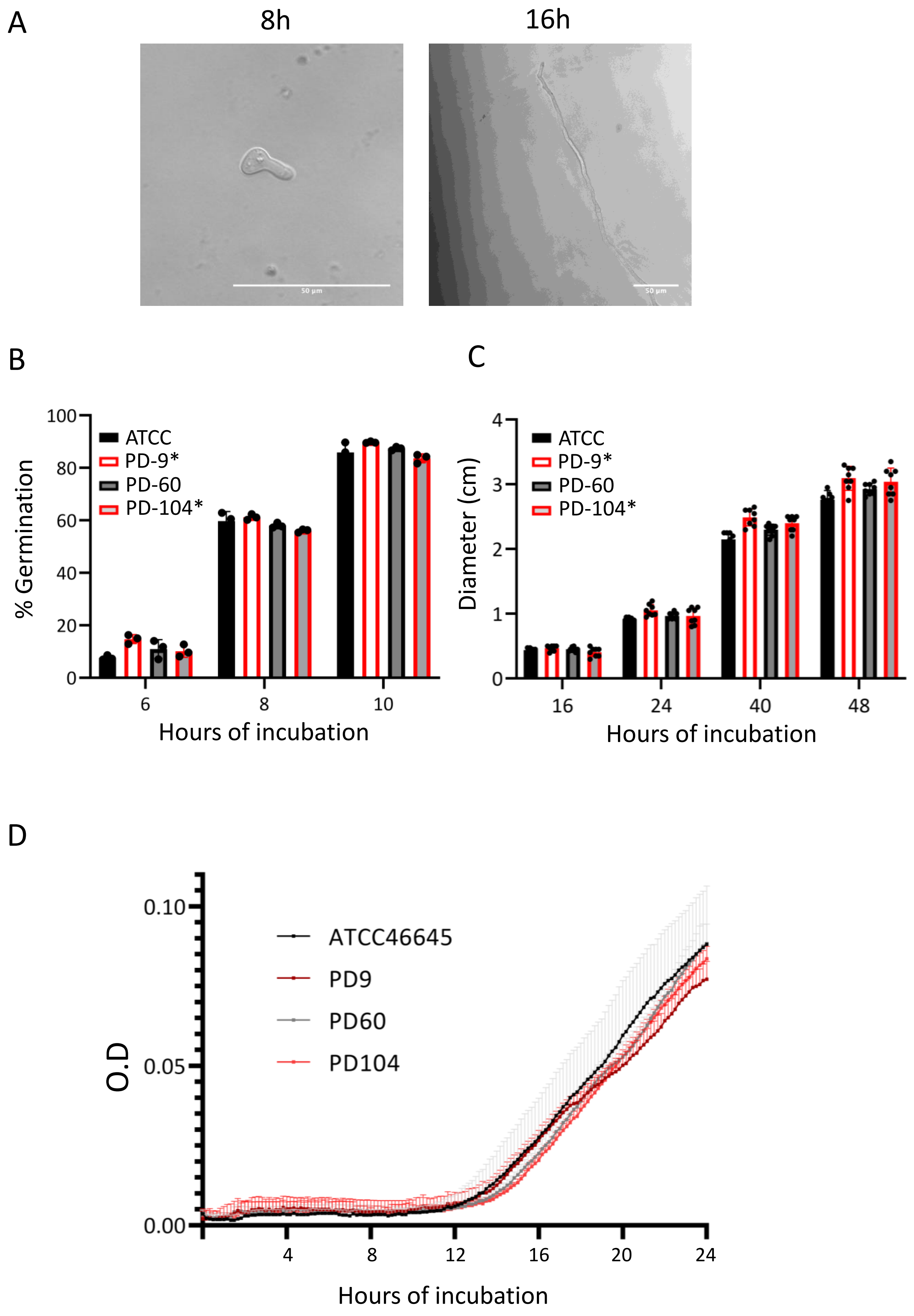

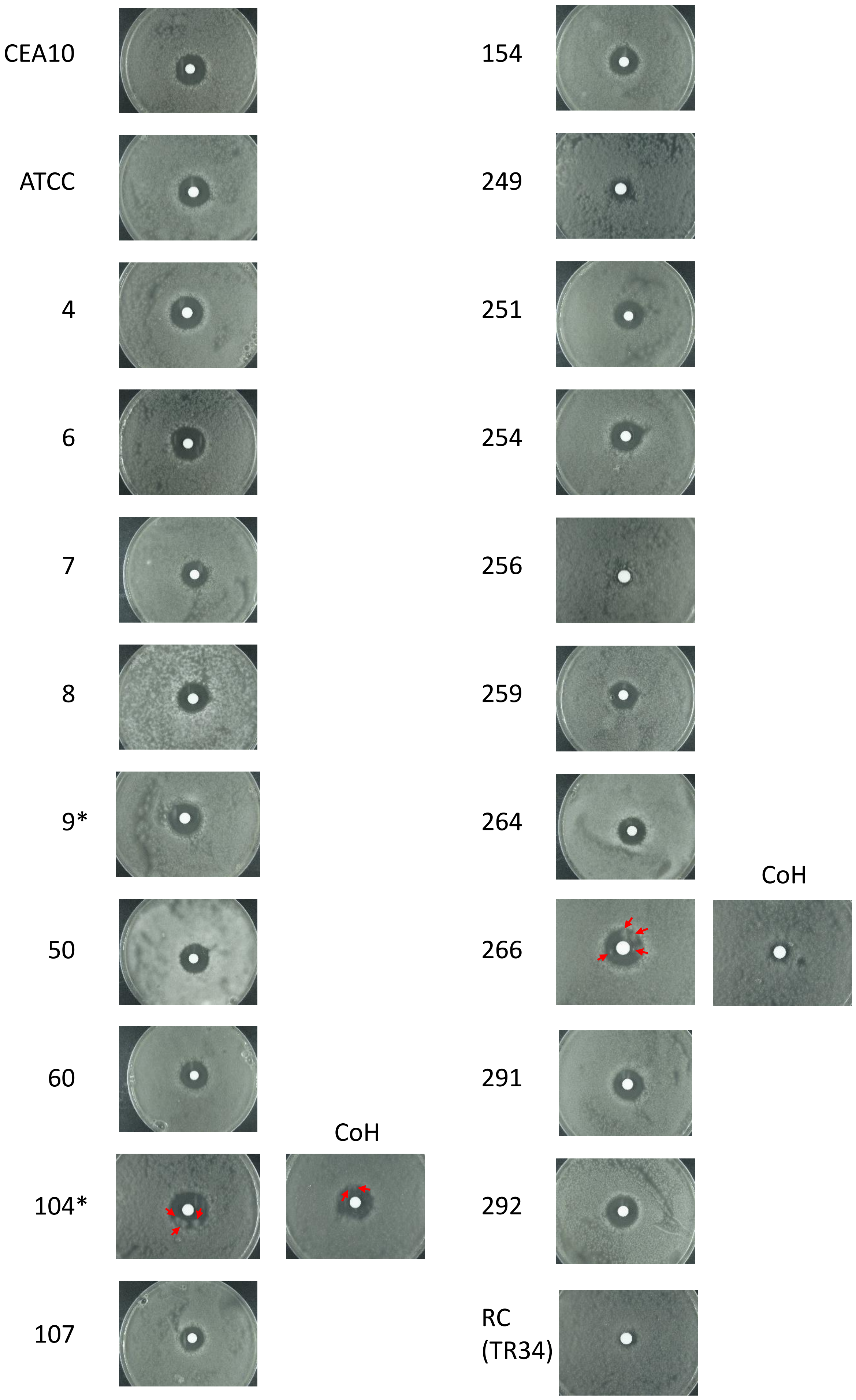

Fig. S7

A

| MIC | Itraconazole (µg/mL) | Isavuconazole (µg/mL) |
|---|---|---|
| ATCC | 0.5 | 1 |
| PD9* | 0.5 | 1 |
| PD60 | 0.5 | 1 |
| PD104 * | 0.5 | 1 |

B

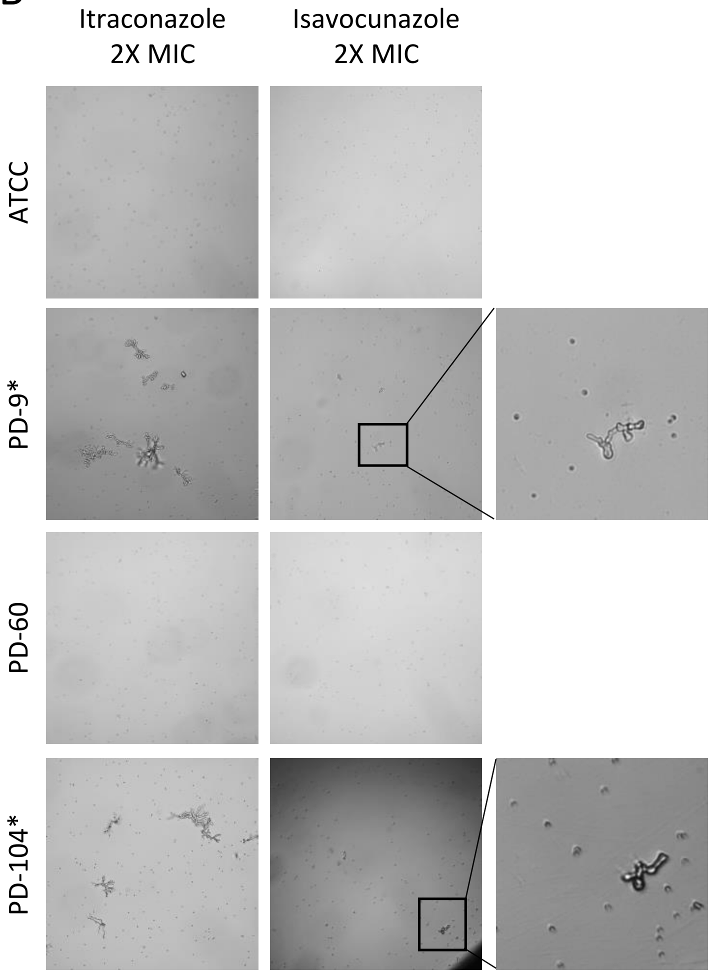


C

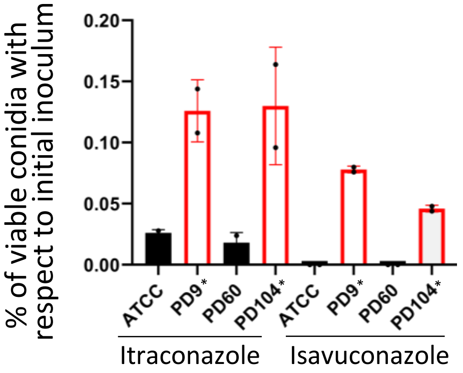

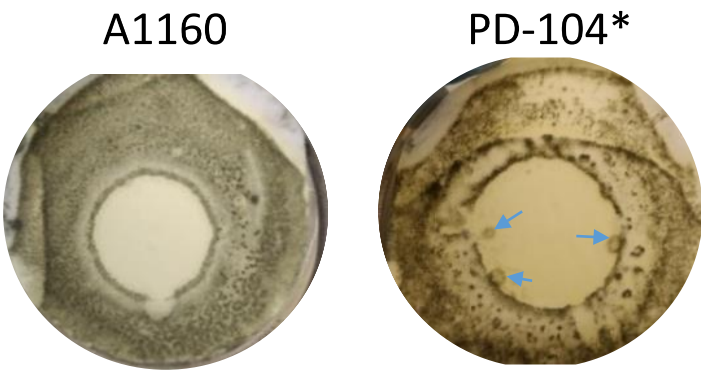

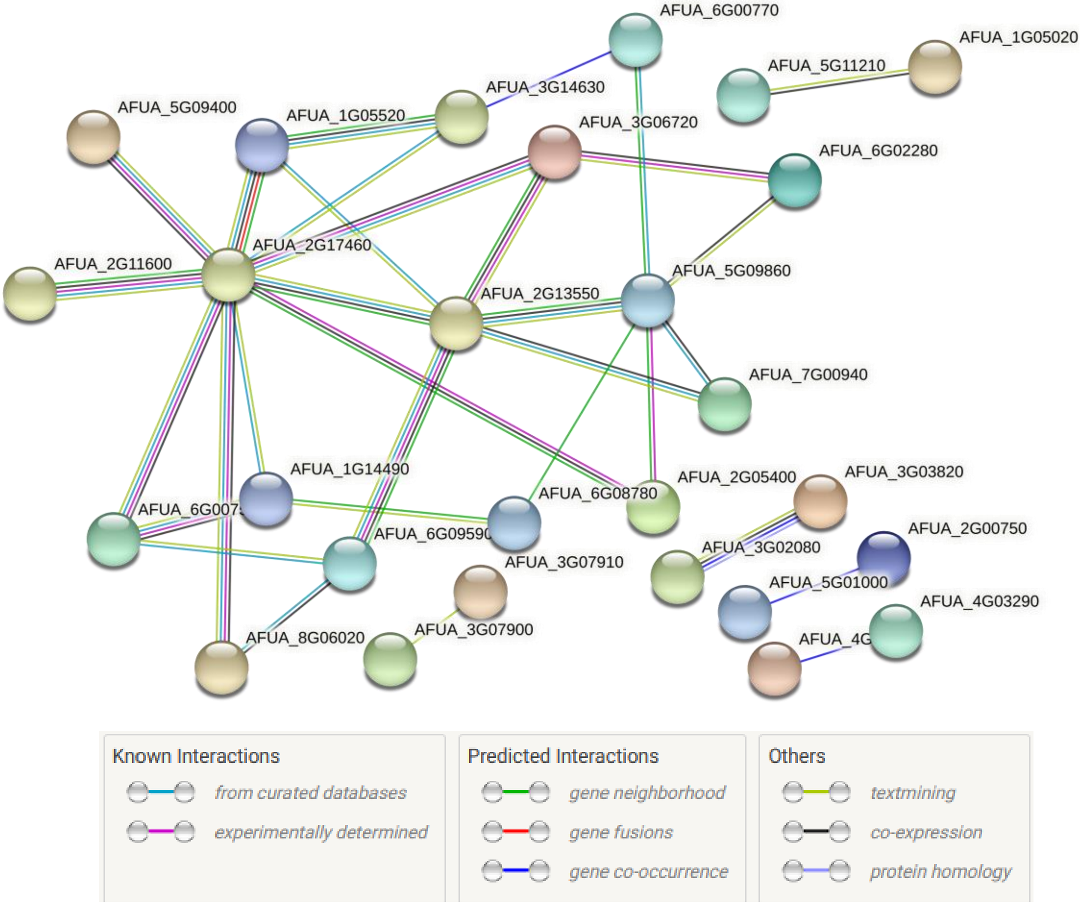

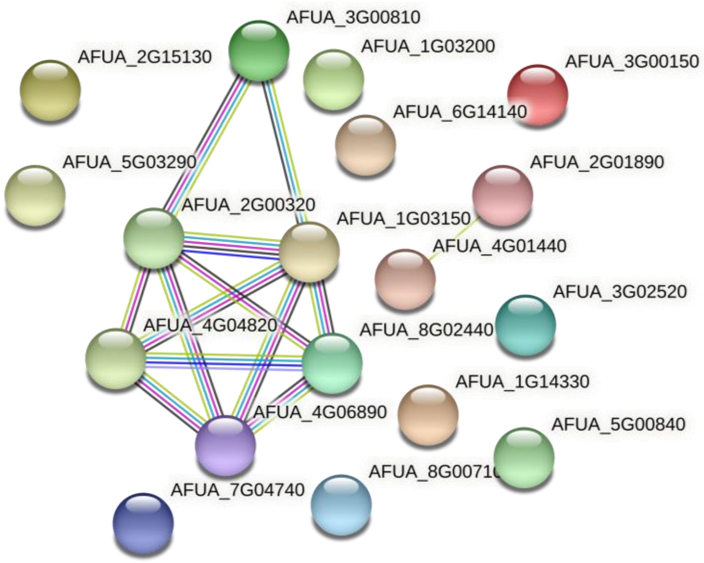

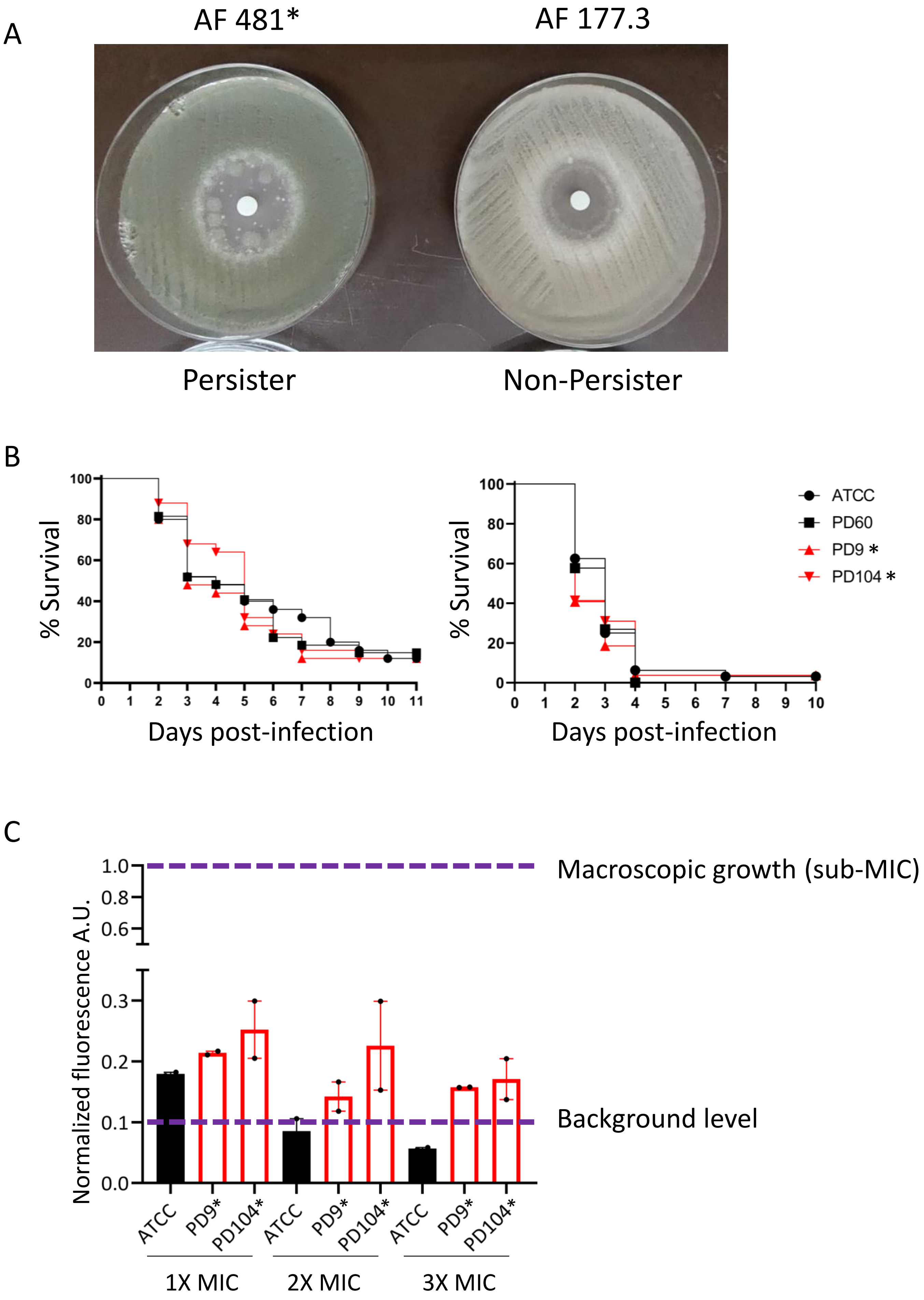
