## Supplementary Table 1 for "*Aspergillus fumigatus* can display persistence to the fungicidal drug voriconazole"

| Isolate | Type | Origin | Colony in halo | Phenotype |
| --- | --- | --- | --- | --- |
| CEA10 | Clinical | Lab | - | Susceptible |
| A1160 | Derivative CEA10 | Lab | - | Susceptible |
| Af293 | Clinical | Lab | - | Susceptible |
| D141 | Clinical | Lab | - | Susceptible |
| ATCC46645 | Clinical | Lab | - | Susceptible |
| RC  TR34/L98H | Resistant control | Lab | + | Resistant |
| PD-4 | Clinical | UK | - | Susceptible |
| PD-6 | Clinical | USA | - | Susceptible |
| PD-8 | Clinical | USA | - | Susceptible |
| PD-249 | Clinical | UK | - | Susceptible |
| PD-251 | Clinical | UK | - | Susceptible |
| PD-254 | Clinical | UK | + | Susceptible  (suspected resistant colony) |
| PD-256 | Clinical | UK | + | Resistant |
| PD-259 | Clinical | UK | -/+ | Possible persister  (low level) |
| PD-264 | Clinical | USA | - | Susceptible |
| PD-266 | Clinical | USA | + | Heteroresistant |
| PD-7 | Environmental | Sweden | - | Susceptible |
| PD-9 | Environmental | USA | + | Persister |
| PD-50 | Environmental | Ireland | - | Susceptible |
| PD-60 | Environmental | Ireland | - | Susceptible |
| PD-104 | Environmental | Thailand | + | Persister |
| PD-107 | Environmental | USA | - | Susceptible |
| PD-154 | Environmental | UK | - | Susceptible |
| PD-291 | Environmental | Thailand | - | Susceptible |
| PD-292 | Environmental | Turkey | - | Susceptible |
