## Supplementary Table 2 for "*Aspergillus fumigatus* can display persistence to the fungicidal drug voriconazole"

| **Isolate** | **Polymorphism** | **Amino acid substitution** |
| --- | --- | --- |
| **ATCC46645** | c822T | S274S |
| **PD-47-9** | t105C, t1182G | S35S, P394P |
| **PD-47-60** | c822T, t1182G | S274S, P394P |
| **PD-47-104** | c822T | S274S |

.
