## Supplementary Table 10 for "*Aspergillus fumigatus* can display persistence to the fungicidal drug voriconazole"

**Quast v5.0.2 RESULTS**:

Assembly                    JA04.contigs (PD-104)

\# contigs (>= 0 bp)         799

\# contigs (>= 1000 bp)      615

\# contigs (>= 5000 bp)      485

\# contigs (>= 10000 bp)     419

\# contigs (>= 25000 bp)     306

\# contigs (>= 50000 bp)     189

Total length (>= 0 bp)      28659847

Total length (>= 1000 bp)   28583670

Total length (>= 5000 bp)   28239428

Total length (>= 10000 bp)  27768809

Total length (>= 25000 bp)  25917775

Total length (>= 50000 bp)  21813374

\# contigs                   661

Largest contig              401152

Total length                28617279

GC (%)                      49.37

N50                         97740

N75                         52625

L50                         85

L75                         183

\# N's per 100 kbp           0.00
